# Structural and Functional Characterization of the KCNJ6 G154C Variant Reveals Severe GIRK2 Channel Gain-of-Function and Opportunities for Drug Repurposing

**DOI:** 10.64898/2026.07.28.741201

**Authors:** Michael A. Netzer, Ilia Steshin, Theres Friesacher, Nathan Dascal, Anna Stary-Weinzinger

## Abstract

G protein-gated inwardly rectifying potassium (GIRK2) channels regulate neuronal excitability and are implicated in neurodevelopmental disorders. A rare KCNJ6 variant, G154C (hGIRK2_G154C_), was identified in a patient with mild Keppen-Lubinsky syndrome features, contrasting with severe phenotypes linked to other selectivity filter mutations. Here we combined molecular dynamics simulations and patch-clamp electrophysiology to characterize the hGIRK2_G154C_ mutant, revealing a widened selectivity filter that resulted in loss of potassium selectivity, aberrant sodium permeation, and loss of inward rectification, indicating a severe gain-of-function phenotype. An in silico and electrophysiological drug screen identified FDA-approved compounds, including nefazodone and eletriptan, that potently inhibited GIRK2 and GIRK2_G154C_ through distinct blocking mechanisms. These findings elucidate the structural and functional impact of the G154C mutation and highlight potential pharmacological tools and therapeutic candidates for the treatment of GIRK2 channelopathies.

## Introduction

G protein-gated inwardly rectifying potassium (GIRK/Kir3) channels are key mediators of inhibitory G protein-coupled receptor signaling and play an essential role in regulating neuronal excitability in the central nervous system (Nguyen et al., 2024).

Among the four mammalian GIRK subunits, GIRK2, encoded by KCNJ6, is highly expressed in the central nervous system and plays important roles in neuronal excitability, motor control, and cognitive function (Hibino et al., 2010; Nguyen et al., 2024).

GIRK channels can assemble as both homo- and heterotetramers. While GIRK1/2 heteromers constitute a major neuronal GIRK population, functional GIRK2 homotetramers have also been described in vivo, including in specific neuronal populations lacking GIRK1 expression (Lüscher and Slesinger, 2010). Importantly, the corresponding murine *weaver* mutation (GIRK2G156S), which affects the homologous residue in the selectivity filter, has been extensively studied in homotetrameric GIRK2 channels and serves as a well-established model for investigating GIRK2 dysfunction (Patil et al., 1995; Kofuji et al., 1996; Slesinger et al., 1996). The extent of protein expression and the level of dysfunction of GIRK1/mGIRK2_G156S_ heterotetramers (containing the *weaver* GIRK2 mutant) remain controversial (Kofuji et al., 1996; Slesinger et al., 1996, 1997). Furthermore, for certain GIRK1/2 stoichiometries, GIRK1 rescues the *weaver* phenotype (Hou et al., 1999). For these reasons, we chose to use the homotetrameric composition of GIRK2 to investigate the primary molecular consequences of the hGIRK2_G154C_ mutation.

In humans, pathogenic variants in KCNJ6 cause Keppen-Lubinsky syndrome (KPLBS), a rare neurodevelopmental disorder characterized by developmental delay, intellectual disability, hypotonia, feeding difficulties, distinctive craniofacial features, and, in many cases, lipodystrophy with an aged appearance. KPLBS was first linked to KCNJ6 through the identification of de novo T152del and G154S variants in affected individuals (Masotti et al., 2015). Subsequent reports expanded the clinical and molecular spectrum of KCNJ6-related disease. A de novo L171R variant, located close to the pore region, was associated with severe hyperkinetic movement disorder and developmental delay, and functional studies showed gain-of-function channel behavior with impaired G-protein activation, loss of K^+^ selectivity, aberrant cation permeability, and constitutive inward current (Horvath et al., 2018). Functional characterization of the selectivity-filter variant G154S further demonstrated profound loss of K^+^ selectivity, aberrant Na^+^ permeation, and altered channel regulation (Friesacher et al., 2022). More recently, additional substitutions of G154 have been reported, including G154R in a child with classical KPLBS features (Gao et al., 2022), highlighting G154 as a recurrent site of pathogenic variation in GIRK2.

Previous studies of the murine *weaver* mutation demonstrated that pathological GIRK2-mediated Na^+^ influx can be pharmacologically suppressed and that inhibition of mutant channel activity rescues granule cell differentiation *in vitro* (Kofuji et al., 1996). These findings suggest that gain-of-function GIRK2 variants may represent tractable therapeutic targets. However, clinically relevant inhibitors of human disease-associated GIRK2 mutants have not been systematically explored.

In contrast to the severe phenotypes associated with G154S and G154R, a de novo G154C variant was reported in an adult patient with mild intellectual disability, subtle dysmorphic features, obsessive–compulsive disorder, and pathological startle responses, representing the mildest KCNJ6-related phenotype described to date (Van Midden et al., 2023). However, despite affecting the same selectivity-filter residue as the severe G154S mutation, the functional consequences of G154C remained unknown.

Here, we investigated the molecular and functional effects of the human GIRK2_G154C_ variant. By combining molecular dynamics simulations, manual and automated patch-clamp electrophysiology, and pharmacological profiling, we aimed to determine how substitution of the selectivity-filter glycine by cysteine alters GIRK2 channel function and to identify pharmacological modulators with potential therapeutic relevance for GIRK2 channelopathies.

## Methods

### Chemicals

Itraconazole (#CAY13288-25) and nefazodone (#CAY10012642-250) were obtained from Szabo-Scandic (Vienna, Austria), verapamil (#V4629-1G) was bought from Sigma-Aldrich (Burlington, USA), and the remaining investigated compounds were purchased from MedChemExpress (South Brunswick, NJ, USA). These included eletriptan (#HY-A0010), lomitapide (#HY-14668), lovastatin (#HY-N0504), maraviroc (#HY-13004), oxacillin (#HY-B0925), permethrin (#HY-B0887), QX-314 (#HY-108505), saquinavir (#HY-17003), Tertiapin-Q (#HY-P1275), and vilazodone (#HY-14261). All chemicals were handled according to suppliers’ suggestions, dissolved in DMSO or distilled water, and stored at −80°C. Tertiapin-Q (TPN-Q) was aliquoted into several 1 mM stocks and underwent only one freeze thaw cycle each to prevent protein degradation. Solutions for pharmacological experiments were always prepared freshly on the day of experiments and DMSO concentrations therein never exceeded 0.1%, except for itraconazole. Due to poor solubility, this stock was prepared at 8 mM, resulting in a final DMSO concentration 0.375% in 30 µM test solution for APC experiments.

### Molecular biology and plasmid constructs

All plasmid constructs used in this study were generated commercially by GenScript Biotech Corporation (Nanjing, China). The human *GNB1* (GenBank accession number NM_002074), *GNG2* (GenBank accession number NM_053064), and *KCNJ6* (GenBank accession number NM_002240) genes were synthesized and cloned into pcDNA3.1(+) mammalian expression vectors. Site-directed mutagenesis was used to generate the KCNJ6 p.Gly154Cys (G154C) expressing plasmid. All sequences and successful mutagenesis were verified by GenScript Biotech Corporation (Nanjing, China) via standard Sanger sequencing. Complete plasmid maps of the three constructs can be found in Supplementary Figure 6. Plasmids were provided in transfection ready midipreps with an endotoxin content of ≤0.01 EU/µg. A standard expression plasmid encoding green fluorescent protein (GFP) was included in all co-transfections.

### Cell culture

Chinese hamster ovary (CHO) – K1 cells (Sigma-Aldrich, Burlington, USA; #8505105) were grown in DMEM/F12 (Thermo Fisher Scientific Inc., Waltham, MA, USA; #10565018) medium supplemented with 10% (v/v) foetal bovine serum (VWR International, Radnor, PA, USA; # HYCLSV30160.03) and 1 mM sodium pyruvate (VWR International, Radnor, PA, USA; #L0642-100), kept in a humidified incubator with 5% CO_2_ and transiently transfected using TurboFect™ Transfection Reagent (Thermo Fisher Scientific Inc., Waltham, MA, USA; #R0531) according to the manufacturer’s protocol. For transfections, 3 µg of target DNA and 0.5 µg of GFP were used with 8 µl of transfection reagent for T-25 flasks. This was scaled up to 9 µg target DNA, 1 µg GFP, and 24 µl reagent for T-75 flasks. Cells transfected with the plasmid carrying the KCNJ6 G154C variant did not survive culturing unless 500 µM QX-314 was co-incubated to block the majority of leak current caused by the mutant channel. Cells were harvested using Accutase® (Sigma-Aldrich, Burlington, USA #A6964) after gentle wash with phosphate buffered saline (Szabo-Scandic, Vienna, Austria; #CTI860015) the day after transfection, replated in 35 mm culture dishes (Corning Inc., Corning, NY, USA; #430165) and used for manual and automated patch clamp recordings 24 – 48h post transfection.

### Manual patch clamp

Extracellular solutions with varying K^+^ concentrations (4, 8, 24, 48, and 96 mM) were used to measure K^+^ conductances. NaCl was equimolarly substituted for KCl so that their combined total concentration remained at 144 mM in each solution. The base extracellular solution contained (in mM): 2 CaCl_2_, 1 MgCl_2_, 10 Glucose, and 10 HEPES. pH was adjusted to 7.4 using NaOH for each extracellular solution. All pharmacological measurements were conducted using 48 mM K^+^ solutions. The intracellular solution contained (in mM): 130 K^+^-gluconate, 20 NaCl, 1 MgCl_2_, 5 HEPES, 1 EGTA, 3 MgATP, and 0.3 Na_2_GTP. To avoid nucleotide breakdown from repeated freeze-thaw cycles, a base solution lacking ATP and GTP was adjusted to pH 7.2 with KOH, divided into 500 µl aliquots, frozen, and stored at – 20°C. On the day of recording, the intracellular solution was finalized by supplementing a freshly thawed base aliquot with pH-adjusted, concentrated frozen stocks of MgATP and Na_2_GTP. Free Mg^2+^ concentration was calculated with CaBuf software (courtesy of Prof. G. Droogmans, Laboratory of Physiology, University of Leuven) to be 547 µM accounting for an ionic strength of 170 mM at 20°C and pH 7.2. Osmolarities were regularly measured with a K-7400S Semi-Micro Osmometer (KNAUER Wissenschaftliche Geräte GmbH, Berlin, Germany) and usually ranged from 290 – 300 mOsm/kg for extracellular solutions, and 280 – 290 mOsm/kg for internal solutions.

A VCPlus-8 Channel Gravity System (ALA Scientific Instruments, Farmingdale, NY, USA) perfusion system was used for all experiments, and liquid flow rate was 0.5 ml/min and controlled on each day of experiments. GFP positive cells were identified using standard fluorescence microscopy with a ZEISS Axio Observer (Zeiss, Oberkochen, Germany) inverted microscope. Glass micropipettes were produced from glass capillaries (Harvard Apparatus, Holliston, MA, USA; #BS4 640792) employing a Sutter P-1000 Micropipette Puller (Sutter Instrument, Novato, CA, USA) and had resistances of 1.5 – 4 MΩ. Data was acquired with a HEKA EPC 10 USB 3.0 Single Patch Clamp Amplifier (HEKA Elektronik GmbH, #895331) at 10 kHz and Bessel filtered at 2.9 kHz using PATCHMASTER NEXT software version 1.5.2 (HEKA Elektronik GmbH, Reutlingen, Germany). Currents were measured with a ramp protocol every 5 s from –120 mV to +60 mV over 0.5 s and cells were held at –20 mV. Ba^2+^ insensitive currents were deducted from GIRK2 wildtype recordings and block was normalized to maximum inward current amplitude. For hGIRK2_G154C_ mutant channels block was normalized to total amplitude, because Ba^2+^ does not block hGIRK2_G154C_ channels and 1 mM QX-314 is insufficient to achieve a complete block of currents. Representative current traces were not adjusted for the liquid junction potential.

### Automated patch clamp

Automated patch clamp experiments were performed using the SyncroPatch 384 (Nanion Technologies GmbH, Munich, Germany) using single-hole NPC-384T S chips, employing the PatchControl384 version 2.3.0 software. Cells were suspended in and added to the chip in divalent-free solution containing (in mM): 140 NaCl, 4 KCl, 5 glucose, and 10 HEPES. A sealing enhancing solution was then added to the cells, which was made of (in mM): 130 NaCl, 4 KCl, 10 CaCl_2_, 1 MgCl_2_, 5 glucose, and 10 HEPES. Cells were then washed twice in standard extracellular recording solution which contained (in mM): 140 NaCl, 4 KCl, 2 CaCl_2_, 1 MgCl_2_, 10 Glucose, and 10 HEPES. High K^+^ extracellular solution #1 contained (in mM): 52 NaCl, 92 KCl, 2 CaCl_2_, 1 MgCl_2_, 10 Glucose, and 10 HEPES. Standard extracellular and high K^+^ solutions were mixed 1:1 on the chip so that the final extracellular K^+^ concentration was 48 mM. Compounds, 400 nM TPN-Q reference blocker, and 3 mM Ba^2+^ containing solutions were respectively diluted at double concentration in High K^+^ extracellular solution #2 which contained (in mM): 96 mM NaCl, 48 mM KCl, 2 CaCl_2_, 1 MgCl_2_, 10 Glucose, and 10 HEPES. All extracellular recording solutions were adjusted to pH 7.4 using NaOH. Intracellular solution was freshly prepared for every experiment and contained (in mM): 80 KF, 50 K^+^-gluconate, 20 NaCl, 1 MgCl_2_, 5 HEPES, 1 EGTA, 3 MgATP, and 0.3 Na_2_GTP. Its pH was adjusted to 7.2 using KOH.

Currents were sampled at 10 kHz and filtered at 3 kHz. All experiments were conducted at room temperature. Quality control parameters were applied to recordings in the following way: cells with a series resistance of >20 MΩ or a capacitance of >40 pF were excluded; minimum membrane resistance thresholds were set at 100 MΩ (in 4 mM K^+^) and 200 MΩ (post TPN-Q application) to account for the basal currents inherent to these conditions. Potassium currents were measured at –140 mV in the whole-cell configuration every 5 s by stepping from the holding potential of –20 mV to –140 mV for 50 ms and then ramping to +60 mV over 200 ms. Block was normalized to maximal TPN-Q inhibition, which is 85 – 90% (Yang, 2013) of GIRK2 mediated currents, and therefore the calculated block could exceed 100%.

### Data analysis and statistics

Manual patch clamp recordings were analyzed using FITMASTER software version 2×92 (HEKA Elektronik GmbH, Reutlingen, Germany), Clampfit version 10.7 (Molecular Devices, San Jose, CA, USA), and SutterPatch version 3.1 (Sutter Instrument, Novato, CA, USA), which was run in IgorPro version 9.05 (WaveMetrics Inc., Portland, OR, USA), which was used to export CSV files for further analysis to control for current rundown using a custom-written script in Python version 3. The script is openly available as detailed in the Data Availability Statement. Automated patch clamp recordings were analyzed using DataControl384 version 3.2.1 (Nanion Technologies GmbH, Munich, Germany), and likewise rundown corrected using a second custom-written script in Python version 3 available as detailed in the Data Availability Statement. Further data and statistical analyses were done using Graphpad Prism version 8 (GraphPad Software, Inc., Boston, MA, USA). Reversal potentials (V_rev_) were determined at the point of intersection of measured currents with the voltage axis. Respective liquid junction potentials were calculated according to the stationary Nernst-Planck equation (Marino et al., 2014) using LJPcalc (RRID:SCR_025044) and subsequently deducted from V_rev_. Inward rectification index (F_ir_) was calculated by dividing the current at 40 mV positive of V_rev_ by the current at 40 mV negative of V_rev_.

The relative permeability ratio of sodium to potassium (pNa/pK, denoted as *r*) was determined according to the Goldman-Hodgkin-Katz voltage equation. To calculate *r* independently of intracellular ion concentrations, we utilized the shift in V_rev_ between two distinct extracellular solutions. Solution 1 contained 8 mM K^+^ (with 136 mM Na^+^) and solution 2 contained 96 mM K^+^ (with 48 mM Na^+^). Experiments were performed at room temperature, assumed to be 20°C, which yields a Nernst slope factor (RT/zF*ln10) of 58.17 mV. The shift in V_rev_ between the two solutions was first used to determine the exponential shift factor, A:

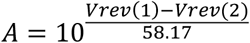

Using the external concentrations ([K^+^]*_o_* and [Na^+^]*_o_*) for solutions 1 and 2, the permeability ratio *r* was subsequently calculated as follows:

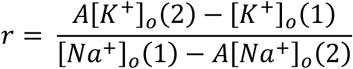

Where [*K*^+^]*_o_*(1) is 8 mM, [*Na*^+^]*_o_*(1) is 136 mM, [*K*^+^]*_o_*(2) is 96 mM, and [*Na*^+^]*_o_*(2) is 48 mM.

Dose-response curves for inhibition of GIRK currents were generated by fitting the normalized current amplitudes to a four-parameter logistic equation (Hill equation) with a variable slope, yielding the IC_50_ and Hill coefficient values.

Voltage-dependence of block of all tested drug concentrations for nefazodone and eletriptan was calculated at voltages from –115 mV to –35 mV. To linearize the voltage dependence, data were transformed and fitted by using simple linear regression:

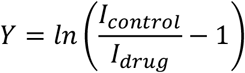

A slope (*m*) significantly different from zero indicated voltage-dependent block. Eletriptan was assumed to be an intracellular pore blocker, because it carries a positive charge (*z* = +1) at physiological pH (Tfelt-Hansen et al., 2000; Milton et al., 2002) and displayed relief of block at more depolarized potentials (see Fig. 7). Woodhull’s electrical distance model (Woodhull, 1973) for voltage-dependent pore block was then used to estimate the relative depth of the eletriptan binding site within the transmembrane electric field. The fractional electrical distance (δ), which represents the fraction of the electric field traversed by the blocker, was calculated as:

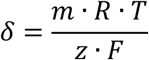

Where *m* is the slope of the linear regression, *R* is the universal gas constant, *T* is the absolute temperature (assumed to be 293.15 K), *z* is the effective charge of the blocker (which is +1 for eletriptan), and *F* is the Faraday constant. The thermodynamic factor *RT/F* was, therefore, approximated as 25.3 mV.

Statistical significance was defined as *p* < 0.05. For permeability ratios and inward rectification indices, unequal variances were found (Brown-Forsythe test, *p* < 0.05), and therefore, this data was analyzed using the Brown-Forsythe and Welch ANOVA followed by the Games-Howell post hoc test. For individual cell Hill slopes and IC_50_ values, equal variances were confirmed (Brown-Forsythe test, *p* > 0.05), allowing for analysis via ordinary one-way ANOVA followed by Tukey’s post hoc test.

### Molecular docking

Docking of FDA-approved compounds was performed using GOLD version 2024.3 (Cambridge Crystallographic Data Centre, Cambridge, UK) (Jones et al., 1997) with the CHEMPLP scoring function (Korb et al., 2009). Ligands were prepared in LigandScout version 4.5 (Inte:Ligand Software-Entwicklungs und Consulting GmbH, Vienna, Austria) (Wolber and Langer, 2005), including generation of protonation states, tautomers, and conformations under default settings. Docking was carried out on GIRK2 structural models (PDB code: 4KFM) using search regions centered on the transmembrane and cytoplasmic pore cavities, with radii chosen to encompass the accessible volume. For each compound, up to 100 poses were generated. Docking solutions were clustered and visually inspected, and compounds were selected for experimental testing based on consistency of binding poses and physicochemical properties, including CNS availability.

### Electrostatic potential calculations

Electrostatic surface potentials were calculated using the pyDelPhi web server (Panday et al., 2026). The GIRK2 cytoplasmic domain structure was converted to PQR format using the PDB2PQR web server (Jurrus et al., 2018) with PROPKA protonation state assignment at pH 7.0 and AMBER force field parameters. Electrostatic potentials were calculated using the default pyDelPhi parameters and mapped onto the molecular surface for visualization in 3Dmol.js.

### Molecular dynamics simulation

Gromacs version 2022.3 (Abraham et al., 2015) was used to perform molecular dynamics simulations with WT and G156C Kir3.2 channels (PDB code: 4KFM; resolution 3.45 Å, Organism: Mus musculus – mGIRK2) (Whorton and MacKinnon, 2013). The protein complex was modeled using the Amber14SB force field (Maier et al., 2015). The channels were embedded in an equilibrated 1-palmitoyl-2-oleoyl-sn-glycero-3-phosphocholine (POPC) membrane, using CHARMM-GUI Membrane Builder (Jo et al., 2008; Wu et al., 2014). Lipids were described using the Lipid21 force field (Dickson et al., 2022). Four phosphatidylinositol 4,5-bisphosphate (PIP_2_) lipids and four Na^+^ ions were positioned according to the crystal structure (Whorton and MacKinnon, 2013). The short-chain PIP_2_ present in the 4KFM crystal structure was replaced with a full-length PIP₂ (18:0/20:4) molecule (1-stearoyl-2-arachidonoyl-sn-glycero-3-phospho-(1D-myo-inositol-4,5-bisphosphate)) as defined in the Lipid21 force field. The phosphate groups were modeled in their fully deprotonated state. K^+^ ions in the selectivity filter were located at positions S0-S4. The system was solvated with TIP3P water molecules (Jorgensen et al., 1983). A total salt concentration of ∼110 mM was added to the simulation system (55 mM KCl, 55 mM NaCl). Ions were modeled using the Joung–Cheatham parameters compatible with the TIP3P water model (Joung and Cheatham, 2008). The C-terminal cysteine of the Gγ subunit was modified with a geranylgeranyl prenylation (CYSG) tail using the CHARMM-GUI Reader & Modeller (Jo et al., 2014; Park et al., 2023). The geranylgeranyl moiety was parameterized within GAFF2 (Wang et al., 2004). Prior to parametrization, the prenylation structure was geometry-optimized, and partial atomic charges were derived using the RESP (Bayly et al., 1993) method based on electrostatic potentials calculated with Gaussian16 (Frisch et al., 2016) at the HF/6-31G* level of theory. The resulting parameters were converted into a GROMACS-compatible topology using ACPYPE (Sousa da Silva and Vranken, 2012).

The system equilibration was carried out in 6 stages: 2 NVT and 4 NPT. During the equilibration process, the protein restriction for the Backbone (BB) and Side chain (SC) was gradually reduced (Supplementary Table 1). For NVT equilibration, a v-rescale thermostat (Bussi et al., 2007) with a coupling constant of 0.1 ps at 310K was used. Both NVT stages lasted 125 ps with an integration time step of 1 fs. For NPT equilibration, a Nose-Hoover thermostat (Evans and Holian, 1985) with a coupling constant of 0.5 ps at 310 K and a semi-isotropic c-rescale barostat (Bernetti and Bussi, 2020) at 1 bar with time constant equal 5 were used. First NPT stage lasted 125 ps with an integration time step of 1 fs. NPT 2-4 stages lasted 500 ps with an integration time step of 2 fs.

For production MD a Nose-Hoover thermostat with a coupling constant of 0.5 ps at 310 K and a semi-isotropic Parrinello–Rahman barostat (Parrinello and Rahman, 1981) at 1 bar with a time constant of 5 ps were used. Long-range electrostatic interactions were calculated every step with the Particle-Mesh Ewald algorithm (Darden et al., 1993) and bonds constrained with the LINCS algorithm (Hess et al., 1997).

Three runs of mGIRK2(WT) and mGIRK2(G156C) were calculated resulting in a total simulation time of 3 μs for the wild-type and for the mutant system. The runs were carried out under electric fields of 300 mV nm^−1^ applied along the z-axis.

Distances, angles, and ion/water flux within the selectivity filter (SF) were analyzed using custom scripts written in Python v3 with the MDAnalysis (Michaud-Agrawal et al., 2011; Gowers et al., 2019), NumPy (Harris et al., 2020), SciPy (Virtanen et al., 2020), and pandas (Reback et al., 2020) libraries. Data visualization was performed using Matplotlib (Hunter, 2007). Ion and water flux through the SF was computed using a grid-based approach with a grid spacing of 3 Å constructed within the SF and adjacent protein cavities. For each grid point, ions and water molecules located within a cutoff distance of 4 Å from the grid center were taken into account.

## Results

### Severe gain of function in the hGIRK2_G154C_ variant

Molecular dynamics (MD) simulations of 1 μs duration were performed for the wild-type (mGIRK2_WT_) and mutant (mGIRK2_G156C_) channels to investigate the impact of the G156C mutation on selectivity filter structure and ion permeation. Ion permeation and water occupancy within the selectivity filter (SF) of mGIRK2_WT_ and mGIRK2_G156C_ over the course of the simulations are shown in Figures 1A,B.

**Figure 1:**
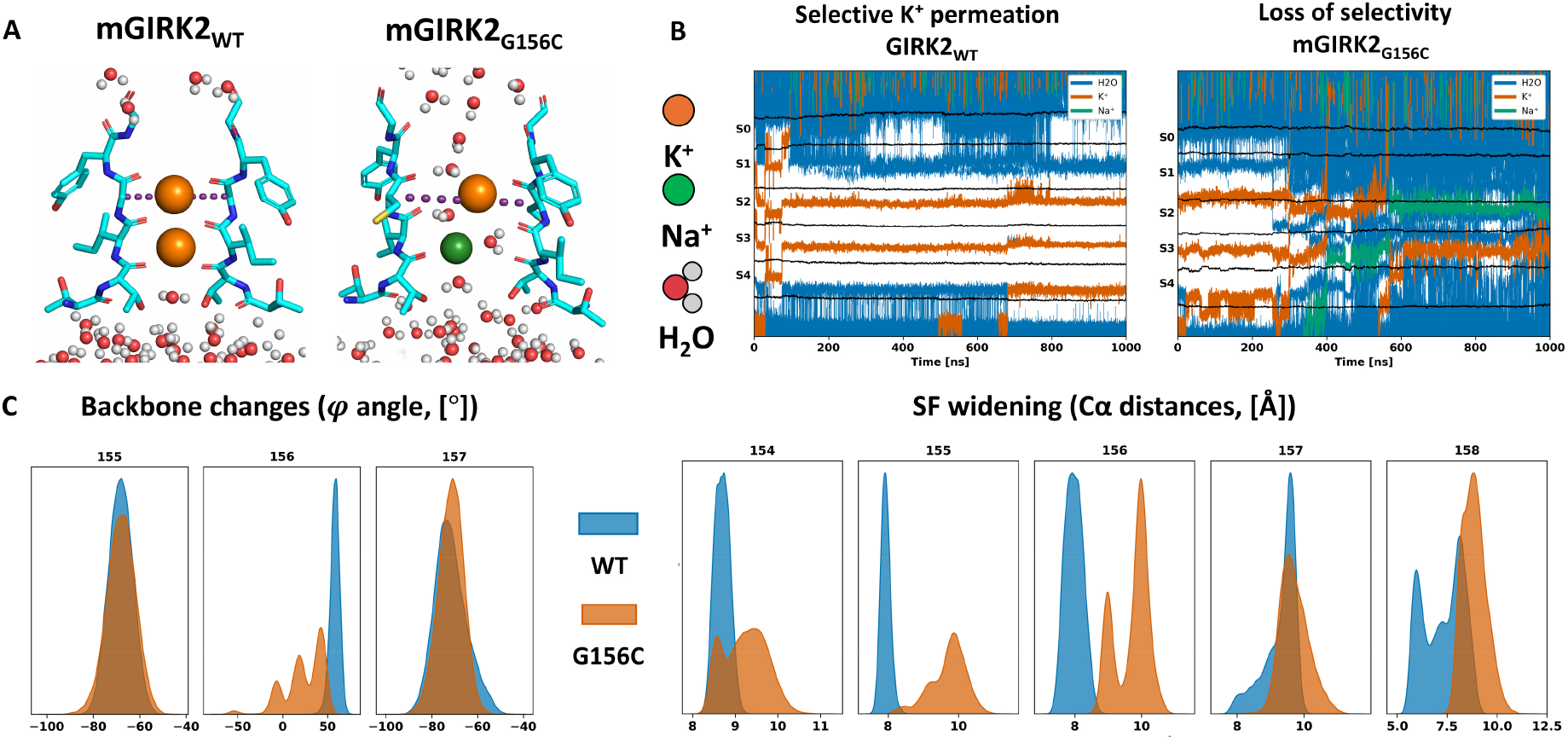
Computational models of mGIRK2_WT_ and mGIRK2_G156C_. **(A)** Molecular dynamics simulations show Na^+^ and H_2_O permeating the selectivity filter (SF) of mGIRK2_G156C_ (the mouse equivalent of the human G154C variant). Widening of the selectivity filter from 7.6 Å in the wild-type to 10.7 Å is observed and indicated by dotted lines. Water molecules and Na^+^ ions can be seen permeating the mutant selectivity filter. **(B)** Na^+^ and water occupancy in the selectivity filter of mGIRK2 and mGIRK2_G156C_ channels over 1 µs of simulation time (sites S0–S4). Data is representative of one of three independent 1 µs simulations conducted at 310 K using the Amber 14SB force field with the channel embedded in an equilibrated POPC membrane. **(C)** Φ-angle distributions of mGIRK2 SF residues (154–158) and distributions of distances between Cα atoms of opposing mGIRK2 SF residues. These metrics quantitatively illustrate the mutation-induced conformational changes and the physical widening of the mutant selectivity filter.

A key difference between the systems is the mutation-induced loss of K^+^ selectivity, as evidenced by Na^+^ permeation and increased water penetration through the SF in mGIRK2_G156C_. Structural analysis revealed that the distances between Cα atoms of SF-lining residues are increased in the mutant compared to the wild type; specifically, for residues 154-158, the average Cα-Cα distance increased by ∼2 Å, from ∼8 Å in the wild type to ∼10 Å in G156C (Figure 1C). Additionally, conformational changes in the backbone dihedral angles (φ, ψ, and ω) were observed for residues 155-157 (Figure 1C, Supplementary Figure 1), disrupting ion coordination by the carbonyl oxygens within the SF. Together, these computational results indicate that the G154C mutation severely compromises SF integrity, resulting in a widened selectivity filter with markedly reduced ion selectivity.

To functionally characterize the consequence of the hGIRK2_G154C_ mutation we heterologously expressed wild-type hGIRK2 and hGIRK2_G154C_ channels in Chinese hamster ovary (CHO) cells and generated current – voltage (I-V) relationships using standard voltage ramps. Strikingly, defining features of GIRK channels such as inward rectification and sensitivity to Ba^2+^ block (Dascal, 1997; Hibino et al., 2010) were completely lost in the hGIRK2_G154C_ mutant channels (Figure 2). Furthermore, hGIRK2_G154C_ mediated currents became susceptible to inhibition by external application of QX-314, a permanently charged derivative of the local anesthetic lidocaine, which was previously shown to block the pore of hGIRK2_G154S_ *weaver* channels (Kofuji et al., 1996; Liss et al., 1999; Slesinger, 2001) (Figure 2A).

**Figure 2:**
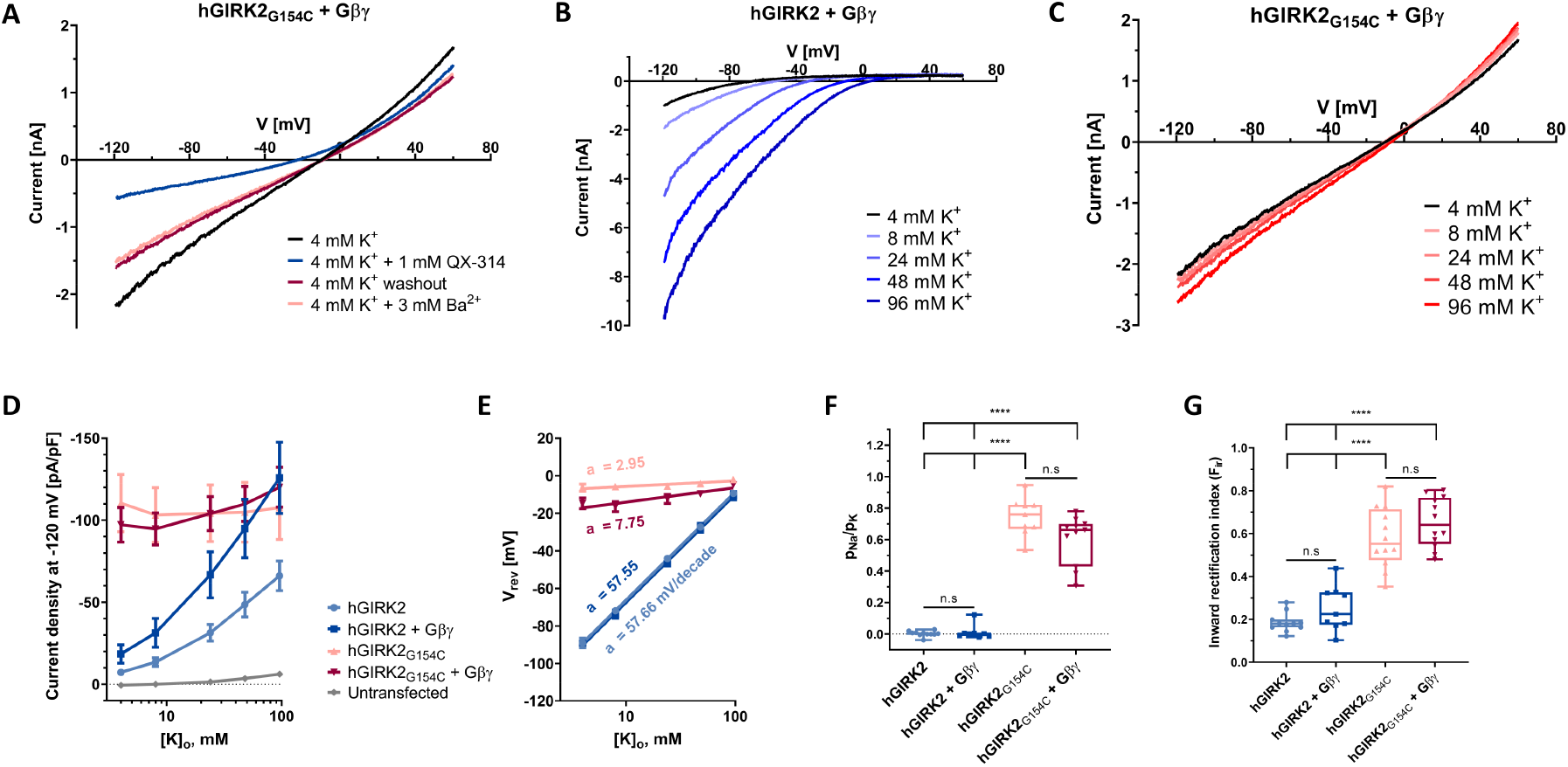
Drastic gain of function and loss of selectivity in the hGIRK2_G154C_ variant. **(A)** Representative manual patch clamp recording in transiently transfected CHO cells showing that hGIRK2_G154C_ mediated currents display loss of inward rectification and that currents can be blocked by QX-314 but not by Ba^2+^. **(B)** and **(C)** Increasing extracellular K^+^ concentrations (equimolarly exchanged with Na^+^) leads to marked increase of inward currents for wildtype channels, but only miniscule increases in current for hGIRK2_G154C_ channels, indicating loss of selectivity. **(D)** and **(E)** Current density and reversal potential (V_rev_) analysis across K^+^ gradients reveal gain of function and an unresponsiveness to co-expressed Gβγ in mutant channels. Error bars represent the standard error of the mean (SEM). The indicated mV/decade values in **(E)** represent the shift in V_rev_ per 10-fold change in external K^+^ concentration. Wild-type channels closely follow Nernstian predictions for a highly K^+^-selective pore (58 mV/decade), whereas mutant channels exhibit marginal shifts (3–8 mV/decade), confirming a profound loss of ion selectivity. **(F)** Relative permeability ratio of Na^+^ vs K^+^ of wild-type and mutant channels as measured by comparing shifts in V_rev_ between 8 mM and 96 mM external K^+^ solutions. **(G)** Inward rectification indices calculated from currents at +40 mV and –40 mV from V_rev_ values in 96 mM K^+^ solutions. Statistical significance in (F) and (G) was determined using the Brown-Forsythe and Welch ANOVA followed by the Games-Howell post hoc test. Significance (*p* < 0.0001) is indicated by four stars (****).

Currents of wildtype and hGIRK2_G154C_ channels were subsequently measured in solutions with increasing concentrations of K^+^, where K^+^ was equimolarly exchanged with Na^+^ to maintain a total monovalent cation concentration of 144 mM (Figures 2B,C). While hGIRK2 wildtype channels showed strong inward rectification, proportionally larger inward currents in higher-concentration K^+^ solutions, and positively shifted reversal potential, V_rev_ (Figure 2B), hGIRK2_G154C_ channels presented as leak currents. The mutant channels showed only minuscule increases in inward currents, a persistent lack of inward rectification across varying K^+^ gradients, and no shift in V_rev_ (Figure 2C), strongly indicating loss of ion selectivity.

Plotting current as a function of external potassium confirmed that wildtype channel currents scale with higher K^+^ concentrations and that co-expression of Gβγ leads to basal activation of hGIRK2 channels (Figure 2D, Supplementary Table 2). This activation was several-fold smaller than previously observed in *Xenopus* oocytes, likely due to the more complex endogenous signaling background of CHO cells (Rubenzik et al., 2001; Hatcher-Solis et al., 2014). Current densities further indicated that hGIRK2_G154C_ channels are insensitive to activation by co-expressed Gβγ, either due to loss of interaction between the mutant channel and Gβγ, as hypothesized for the hGIRK2_G154S_ mutation (Navarro et al., 1996; Friesacher et al., 2022), or because the structurally deformed hGIRK2_G154C_ channels are already in a maximally activated state (Figure 2D, Supplementary Table 2).

Reversal potentials (V_rev_) were calculated and plotted against external K^+^ (Figure 2E, Supplementary Table 3). Wildtype channels followed predictions by the Nernst equation for a highly selective potassium channel (linear relation between V_rev_ and K^+^ concentration; the slope was 57 mV per 10-fold change in K^+^, Figure 2E). In contrast, the hGIRK_G154C_ mutant exhibited only marginal changes in V_rev_ across the tested solutions (3 – 8 mV per 10-fold change in K^+^, Figure 2E), further supporting that K^+^ is not the sole charge carrier in hGIRK2_G154C_ mediated currents. Relative permeability ratios of Na^+^ to K^+^ (pNa/pK) were calculated using the Goldman-Hodgkin-Katz equation and revealed drastically increased sodium permeation in hGIRK_G154C_ channels, which was not observed in wildtype channels, confirming our computational findings (Figure 2F, Supplementary Tables 4,5).

Finally, inward rectification ratios were calculated for hGIRK2 (0.186 ± 0.013) and hGIRK2 with Gβγ co-expressed (0.252 ± 0.035). These values indicated strong inward rectification, which is in line with previous reports (Figure 2G, Supplementary Tables 6,7) (Friesacher et al., 2022). Significantly higher inward rectification ratios were found for hGIRK2_G154C_ channels (0.580 ± 0.040) and hGIRK2_G154C_ channels with Gβγ co-expressed (0.652 ± 0.034), confirming loss of rectification by voltage dependent polyamine block (Figure 2G, Supplementary Tables 6,7). Together, these data demonstrate a gain of function and loss of ion selectivity in the homotetrameric hGIRK2_G154C_ channel.

### In silico screening of FDA-approved drugs identifies potential hGIRK2_G154C_ pore blockers

Given the pronounced gain-of-function of the G154C variant and the lack of targeted therapeutic options for GIRK2 channelopathies, we used docking as a structure-guided prioritization step to select clinically approved compounds for functional testing. FDA-approved, BBB-permeable compounds were docked with GOLD into the transmembrane and cytoplasmic domain cavities of mGIRK2. Docking poses were inspected and clustered, and a limited set of candidates was prioritized for experimental validation based on pose plausibility, physicochemical properties (including CNS availability) and expert assessment of pharmacological and translational relevance. This set included verapamil, previously reported to inhibit the *weaver* GIRK2 mutant in *Xenopus* oocytes and rescue *weaver* granule cells (Kofuji et al., 1996).

### Automated patch clamp drug repurposing screen

To functionally validate the *in silico* selected hits, we evaluated the 11 prioritized compounds at a concentration of 30 µM against the hGIRK2 wildtype channel with co-expressed Gβγ using the high-throughput automated patch clamp (APC) platform SyncroPatch 384 (Nanion Technologies GmbH) (Figure 3A, Supplementary Table 8). Direct testing against the hGIRK2_G154C_ channel on the APC platform proved unfeasible, as overexpression of the mutant channel induced significant cytotoxicity that was only partially rescued by co-incubation with QX-314 during culturing. Maximal inward currents were measured at –140 mV in 48 mM K^+^ solutions prior to the addition of 30 µM compounds, followed sequentially by 400 nM Tertiapin-Q, and 3 mM Ba^2+^ (Figure 3B shows a representative recording of verapamil, Supplementary Figures 5A-K show representatives of the remaining compounds).

**Figure 3:**
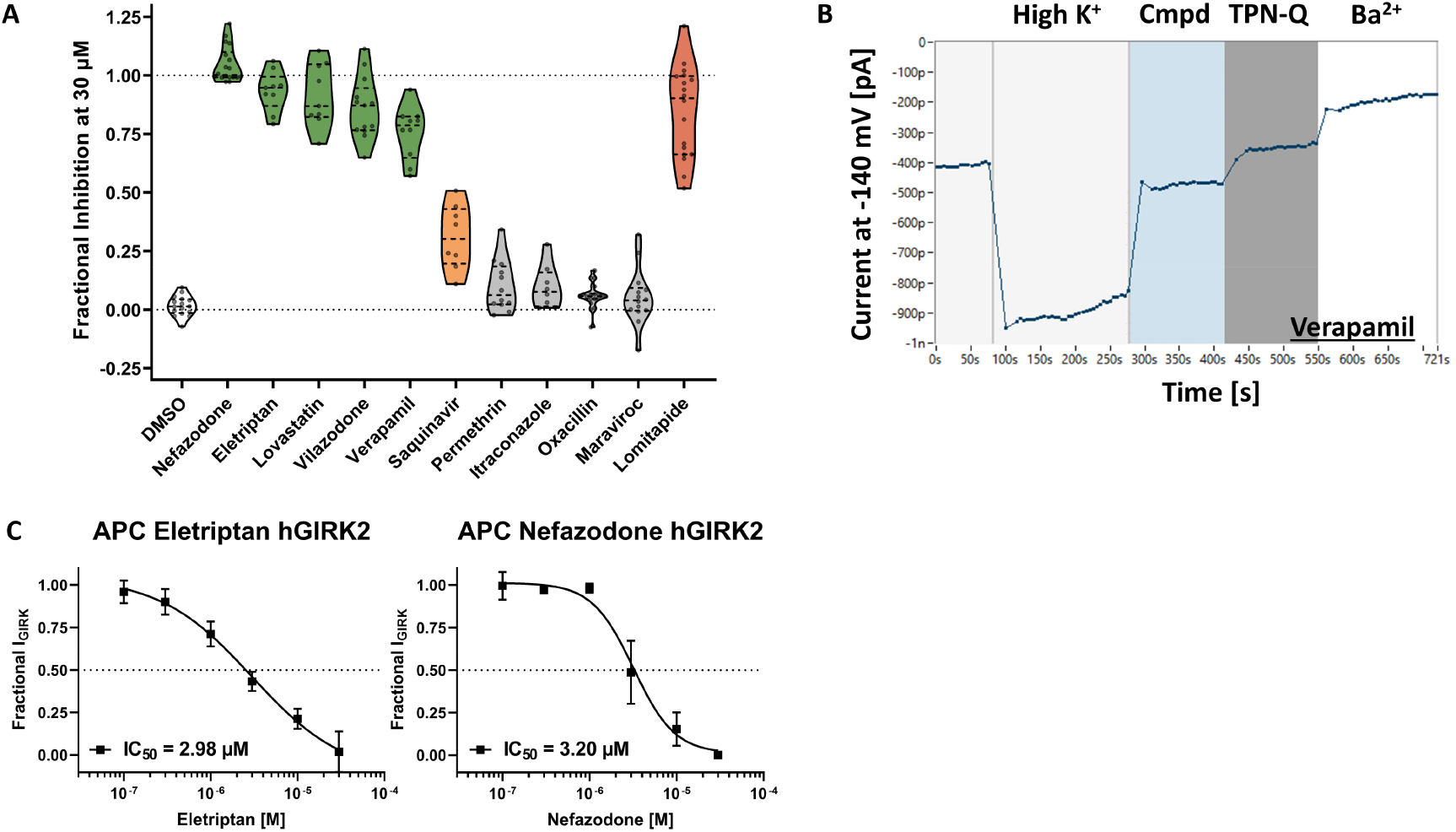
Automated patch clamp screen identifies four hGIRK2 inhibitors. **(A)** Fractional inhibition of hGIRK2 + Gβγ wildtype currents by 11 selected FDA-approved compounds at 30 µM concentration. Data was acquired utilizing the automated patch clamp (APC) platform SyncroPatch 384. Inhibition is normalized to a 400 nM TPN-Q reference block; because TPN-Q inhibits ∼85–90% of total GIRK2 current, calculated fractional inhibition can exceed 1.0. Compounds in green show >70% inhibition, compounds in yellow show 30 – 70% inhibition, compounds in grey show <30% inhibition. Lomitapide appears as a false positive due to acute non-specific membrane destabilization, indicated by a drastic drop in cellular capacitance, and is colored red. Violin plots denote the probability density and distribution of fractional inhibition across individual cells, and internal lines represent the median and interquartile ranges. **(B)** Assay trace example showing a cell treated with 30 µM verapamil (positive control), with 400 nM TPN-Q serving as the reference blocker for normalization. Block with 3 mM Ba^2+^ was used for quality control at the end of the assay. Data points show mean currents of cells which passed quality control criteria from one representative run. **(C)** Dose-response curves for eletriptan and nefazodone generated via APC, fitted to a four-parameter logistic (Hill) equation. Error bars denote SD.

Five compounds significantly inhibited (>70% inhibition) hGIRK2 currents: the structurally related antidepressants nefazodone and vilazodone; the abortive migraine drug eletriptan; the cholesterol-lowering statin lovastatin; and the Ca^2+^ channel blocker verapamil, which served as positive control. We generated dose-response curves for nefazodone and eletriptan and determined IC_50_ values of 2.98 µM for eletriptan and of 3.20 µM for nefazodone (Figure 3C, Supplementary Tables 9,10). As we worked with transient transfections, the success rate of the APC assay was not high enough to satisfactorily resolve the precise mechanisms of action of nefazodone and eletriptan, therefore, we transitioned to manual patch clamp (MPC) for detailed dose-response and biophysical analysis.

### Nefazodone acts as a voltage-independent, cooperative blocker

Dose-response curves were generated for nefazodone (Figure 4A-G) in the presence and absence of co-expressed Gβγ, for both wildtype hGIRK2 and hGIRK2_G154C_ mutant channels using MPC (Figures 4B,C). Wildtype IC_50_ values were 5.12 µM in hGIRK2 and 2.30 µM in hGIRK2 + Gβγ, matching our APC data (Supplementary Tables 11,12). Strikingly, wildtype channels exhibited Hill slopes of 1.22 (basal) and 3.13 (+ Gβγ), indicating potential positive cooperative block requiring multiple drug molecules. To ensure this steep Hill slope was not an artifact of pooling data, we evaluated the dose-response fits for individual cells. This cell-by-cell analysis demonstrated that both the Hill slopes and IC_50_ values remained consistent across individual recordings, validating the cooperativity observed in the averaged data (Supplementary Figures 3,4; Supplementary Tables 13,14). While concentrations up to 1 µM had negligible effects, 3 µM and higher yielded potent, rapid current reductions (Figures 4B,D,F).

**Figure 4:**
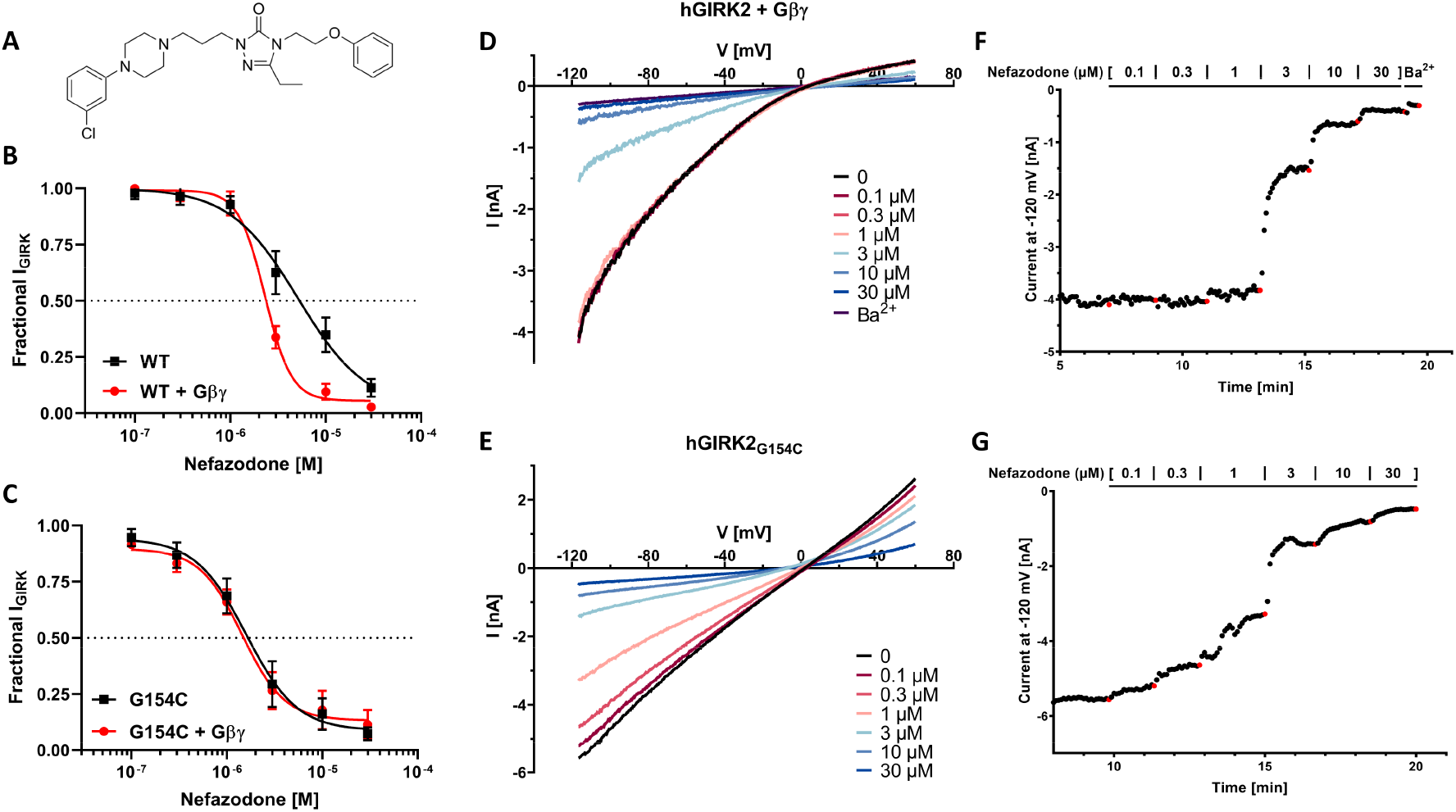
Nefazodone pharmacology using manual patch clamp. **(A)** Chemical structure of nefazodone, sourced from Wikimedia Commons (Public Domain). **(B)** and **(C)** Dose-response curves of wild-type hGIRK2 and mutant G154C channels in the presence and absence of co-expressed Gβγ. Currents were measured at –120 mV in 48 mM external K^+^ solutions and normalized to 3 mM Ba^2+^ block for wild-type conditions or maximum negative amplitude for mutant conditions. Error bars denote SD. **(D)** and **(E)** Representative current traces for wild-type and mutant channels. **(F)** and **(G)** Corresponding currents over time from the respective representative traces in (D) and (E).

In wild-type channels, fractional block at sub-µM nefazodone concentrations is virtually absent. We propose that this sharp activation threshold is the result of the high cooperativity of the interaction with the channel. The steep Hill coefficient of 3.13 compresses the dose-response transition into a highly restricted concentration window. Conversely, the mutant hGIRK2_G154C_ channels yielded lower IC_50_ values of 1.60 µM and 1.45 µM in the presence and absence of co-expressed Gβγ, respectively, along with shallower Hill slopes of 1.63 and 1.95 when compared to the wildtype GIRK2 + Gβγ condition. By reducing the cooperative threshold, the range of the dose-response curve gets broadened, enabling robust and detectable block at lower nefazodone concentrations in the mutant channel (Figures 4C,E,G).

Voltage dependence of block was evaluated from –115 to –35 mV (Figures 5A-D); as only 1 of 24 slopes significantly deviated from zero (Supplementary Table 15), we conclude that nefazodone block of both wildtype and mutant channels is voltage-independent.

**Figure 5:**
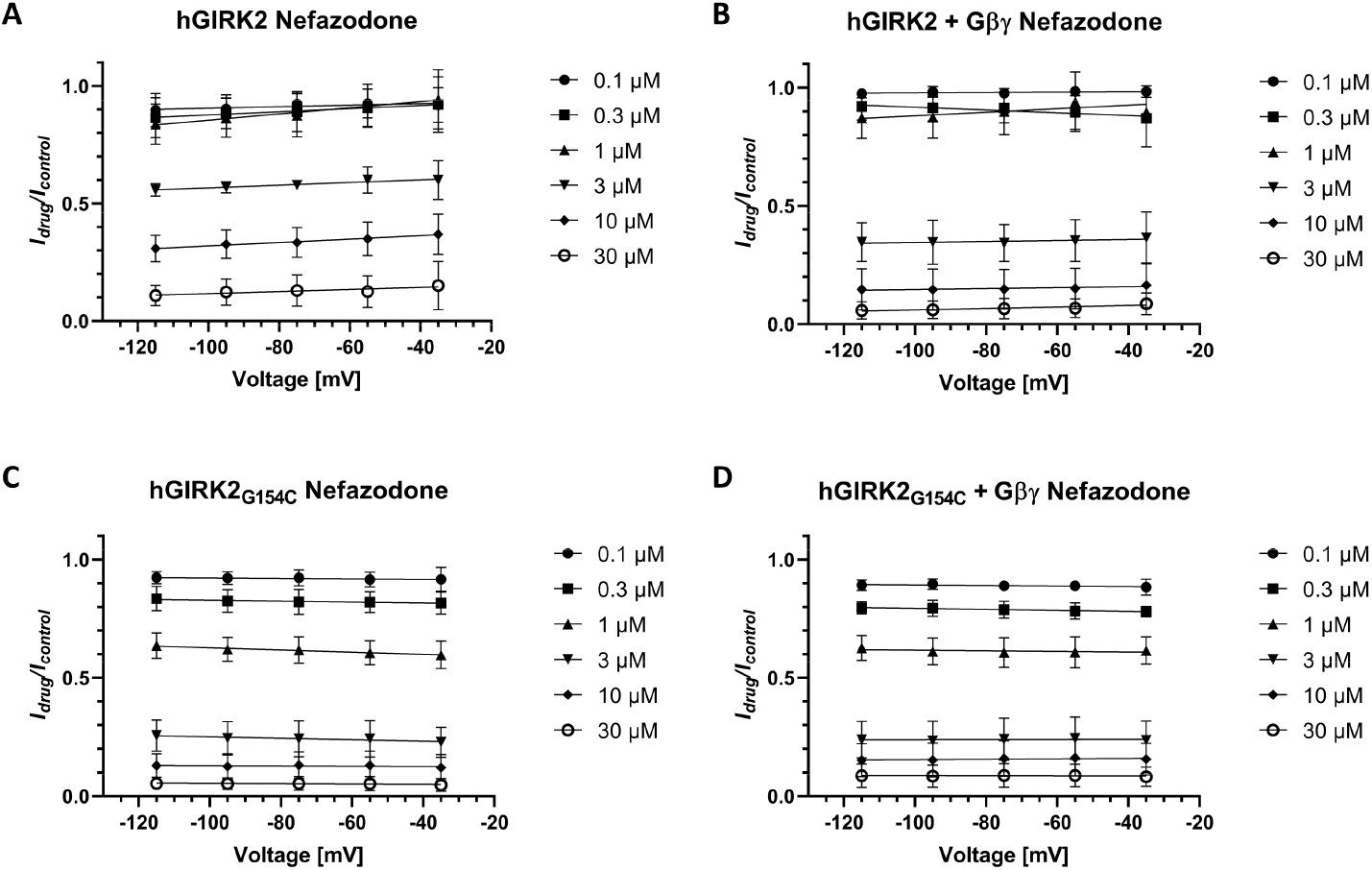
Nefazodone block is voltage independent. **(A) – (D)** Plots showing current inhibition of tested drug concentrations of nefazodone (0.1 to 30 µM) for wild-type and mutant channels in presence and absence of Gβγ across test voltages from –115 mV to –35 mV. Data were transformed and evaluated using simple linear regression. Slopes did not significantly deviate from zero (*p* > 0.05) across nearly all conditions (see Supplementary Table 15), indicating strictly voltage-independent channel block.

### Eletriptan acts as a voltage dependent, intracellular pore blocker

We performed an analogous MPC dose-response analysis for eletriptan (Figures 6A-G). Wildtype hGIRK2 currents were inhibited with IC_50_ values of 2.88 µM and 1.49 µM, with Hill slopes of 0.77 and 0.78, in the presence and absence of Gβγ, respectively (Figures 6B,D,F; Supplementary Tables 16,17), and MPC data closely matched APC data (Figures 3C, 6B,C). In wild-type channels, co-expression of Gβγ induced a left-shift in the dose-response curve, increasing eletriptan potency (Figures 6B,C). Remarkably, this Gβγ-dependent left-shift was lost in the hGIRK2_G154C_ mutant. Mutant hGIRK2_G154C_ channel currents were similarly inhibited with IC_50_ values of 3.09 µM and 3.68 µM, with Hill slopes of 0.84 and 1.07 (Figures 6C,E,G; Supplementary Tables 16,17).

**Figure 6:**
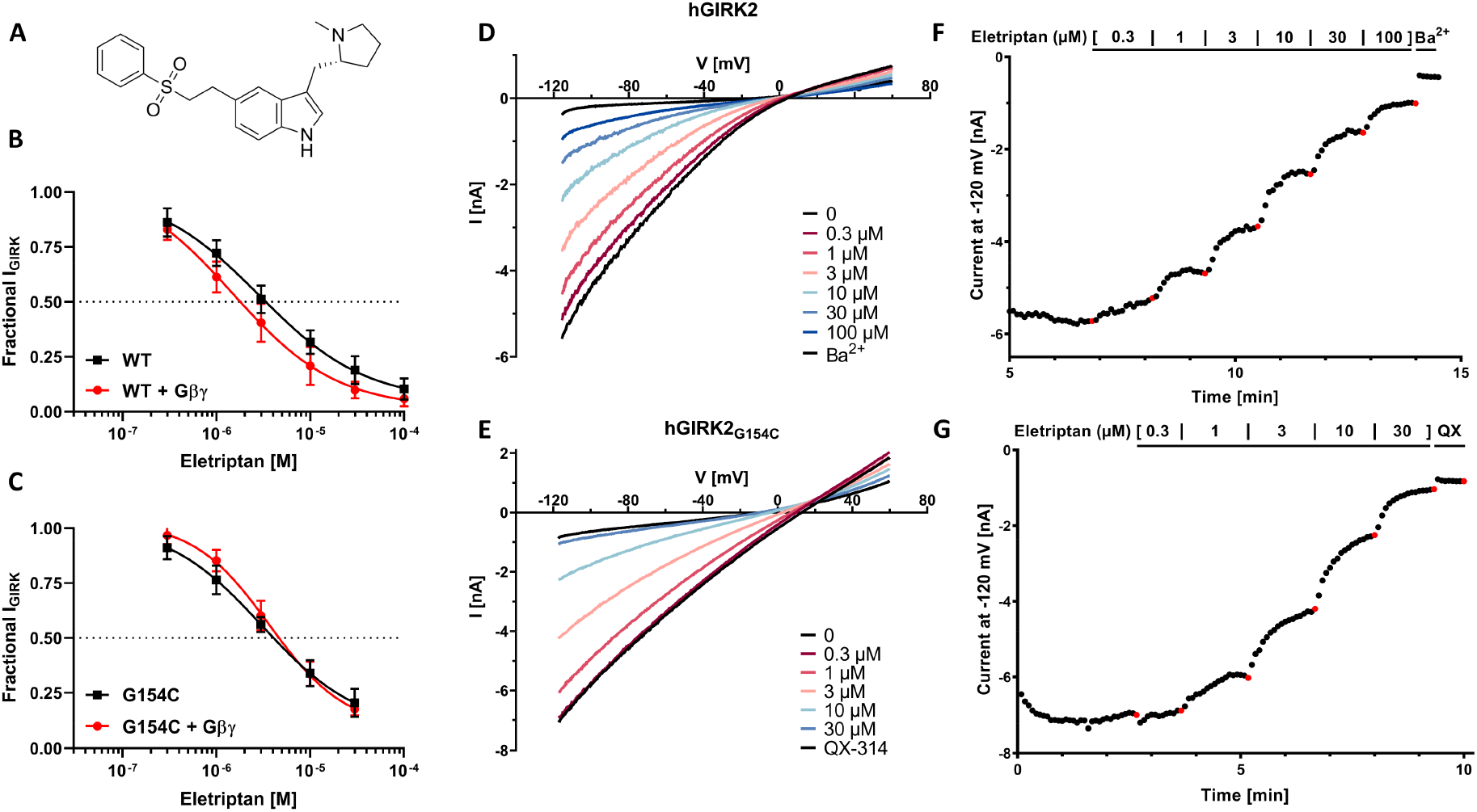
Eletriptan pharmacology using manual patch clamp. **(A)** Chemical structure of eletriptan, sourced from Wikimedia Commons (Public Domain). **(B)** and **(C)** Dose-response curves of wild-type hGIRK2 and mutant G154C channels in presence and absence of Gβγ. Currents were measured at – 120 mV in 48 mM external K^+^ solutions and normalized to 3 mM Ba^2+^ block for wild-type conditions or to 1 mM QX-314 block for mutant conditions. Error bars denote SD. **(D)** and **(E)** Representative current traces for wild-type and mutant channels. **(F)** and **(G)** Corresponding currents over time from the respective traces in (D) and (E).

In contrast to nefazodone, the voltage dependence of eletriptan block was highly significant (Figures 7A-D). Most slopes significantly deviated from zero (Supplementary Table 18), revealing increasingly potent block at more hyperpolarized potentials. Assuming a simple, noncooperative interaction, the depth of the eletriptan binding site within the pore was estimated using Woodhull’s electrical distance model (Woodhull, 1973). The fractional electrical distance (δ) was calculated to be 0.27, 0.29, 0.24, and 0.21 for the conditions hGIRK2, hGIRK2 + Gβγ, hGIRK2_G154C_, and hGIRK2_G154C_ + Gβγ, respectively.

**Figure 7:**
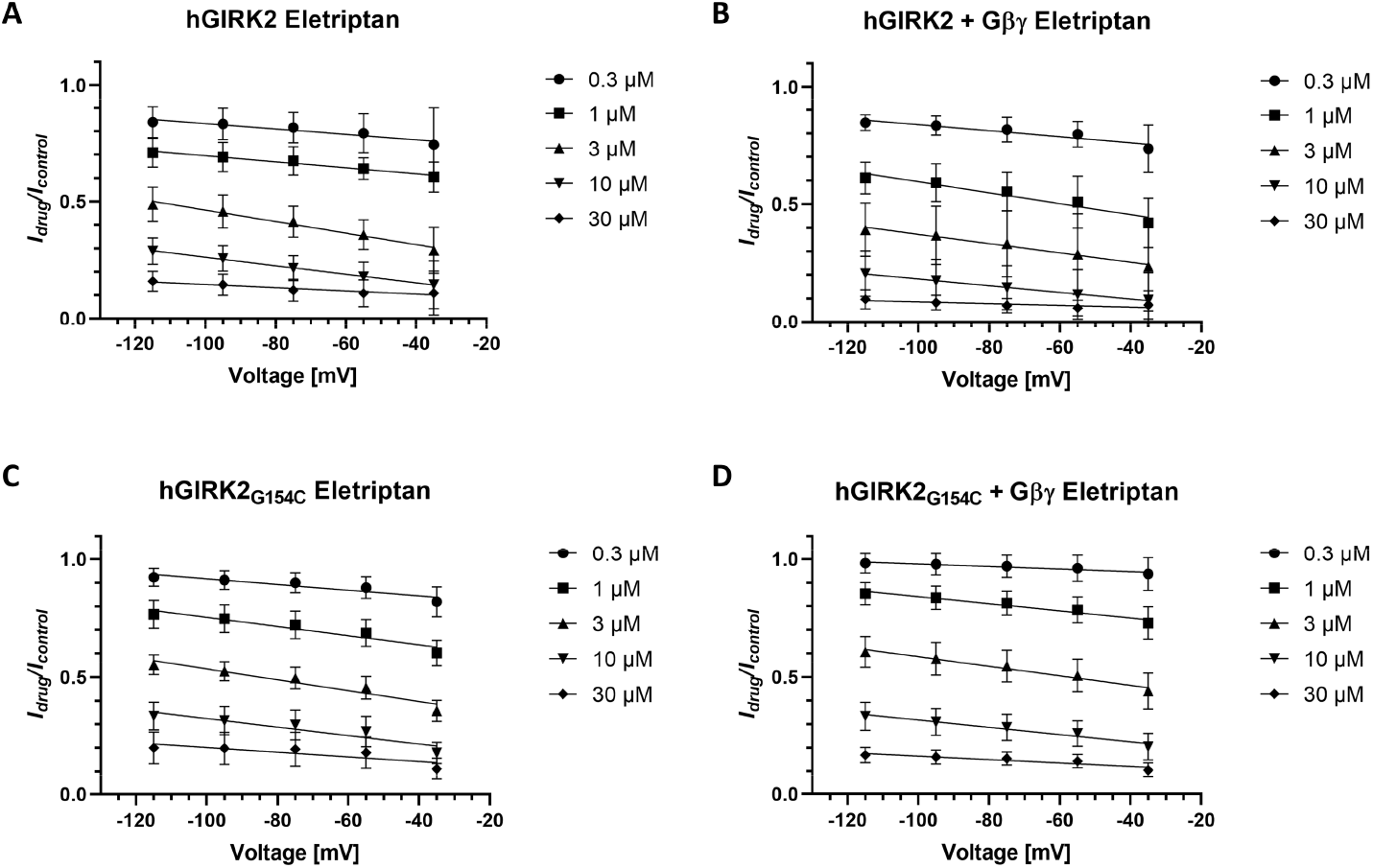
Eletriptan block is voltage dependent. **(A) – (D)** Plots showing current inhibition of tested drug concentrations of eletriptan (0.3 to 30 µM) for wild-type and mutant channels in presence and absence of Gβγ across test voltages from –115 mV to –35 mV. Data were evaluated using simple linear regression. Slopes significantly deviated from zero for multiple concentrations across all channel conditions (see Supplementary Table 18), indicating increasingly potent block at hyperpolarized potentials consistent with an intracellular pore-blocking mechanism.

To assess whether the voltage-dependent inhibition by eletriptan was compatible with its predicted binding location, we revisited the docking poses obtained during the initial *in silico* screen. A representative high-ranked pose localized eletriptan to the central cavity of the GIRK2 cytoplasmic domain below the G-loop gate, where the positively charged ligand is positioned within a strongly electronegative intracellular vestibule formed by residues from multiple subunits (Figure 8). This location is consistent with the shallow electrical distance obtained from Woodhull analysis and supports an intracellular pore-blocking mechanism.

**Figure 8:**
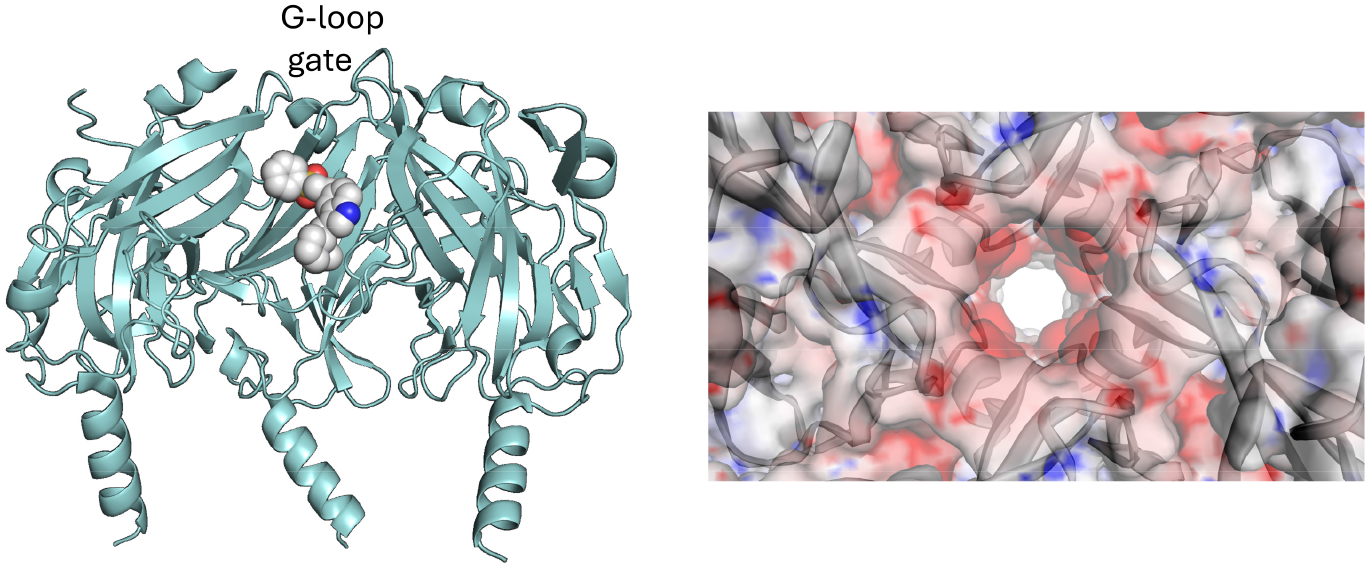
Predicted docking pose of eletriptan in GIRK2. Molecular docking of eletriptan was performed using GOLD with the ChemPLP scoring function. The channel is shown in cartoon representation and the ligand as spheres. Left: Representative docking pose of eletriptan within the central cavity of the GIRK2 cytoplasmic domain below the G-loop gate. For clarity, the front subunit is omitted. Right: Electrostatic surface potential of the GIRK2 cytoplasmic vestibule viewed from the intracellular side toward the G-loop gate. Red indicates negative electrostatic potential, and blue indicates positive electrostatic potential. The predicted binding site is located within a strongly electronegative intracellular vestibule, physically occluding the ion pathway just outside the Helix Bundle Crossing (HBC) gate.

## Discussion

We previously reported that the Keppen-Lubinsky-syndrome (KPLBS)-causing mutation hGIRK2_G154S_ leads to altered SF geometry, aberrant ion selectivity, and structural changes in the Gβγ binding site using a combination of electrophysiological methods and computational models of the murine mGIRK2_G156S_ homolog (Friesacher et al., 2022). Here we investigate the effects of the G154C mutation on hGIRK2 homomer structure and function. Simulations revealed widening of the SF by approximately 3 Å, accompanied by loss of K^+^ selectivity, increased water penetration, and Na^+^ permeation through the SF. These computational findings were independently confirmed by electrophysiological characterization, which further demonstrated loss of inward rectification in hGIRK2_G154C_ mutant channels. Furthermore, we demonstrated loss of block by Ba^2+^ and the appearance of block by QX-314. The hGIRK2_G154C_ homomeric mutant channel is, therefore, remarkably similar to the hGIRK2_G154S_ homomeric mutant channel in biophysical properties.

The close similarity between the G154C and G154S variants indicates that the observed channel dysfunction is not driven by the specific chemical properties of the substituted residue. Instead, our data suggest that Gly154 plays a critical structural role in maintaining selectivity filter architecture. Replacement of this conserved glycine by either cysteine or serine produces a highly similar biophysical phenotype, despite the distinct side-chain properties of these amino acids. Similar loss-of-selectivity phenotypes have been reported for mutations affecting the homologous selectivity-filter glycine in another Kir3 subunit, GIRK4/KCNJ5 (Scholl et al., 2012), suggesting a conserved structural role of this residue across the GIRK channel family.

These results are difficult to reconcile with the phenotype of the single identified patient harboring the G154C mutation. Clinically, this patient exhibited a relatively mild symptom profile in contrast to the severe phenotypes presenting in other known KPLBS patients (De Brasi et al., 2003; Basel-Vanagaite et al., 2009; Masotti et al., 2015; Horvath et al., 2018; Van Midden et al., 2023). The considerable phenotypic variability observed among patients carrying KCNJ6 variants suggests that channel dysfunction alone may not fully determine disease severity. Differences in genetic background, modifier variants, developmental compensation, and environmental influences may all contribute to clinical outcome. As seen with the well characterized mGIRK2*_WV_* mutation (G156S, corresponding to the human G154S), where the mGIRK1_WT_ subunits can “rescue” the wild-type phenotype (Hou et al., 1999), similar compensatory interactions with other channel subunits might mitigate the severe biophysical defects of the hGIRK2_G154C_ variant *in vivo*. Alternatively, somatic mosaicism cannot be excluded, as the G154C variant was identified using peripheral blood DNA and may not reflect the mutational burden across all tissues (Thorpe et al., 2020; Van Midden et al., 2023).

Given the severity of the identified biophysical defects and the current absence of targeted therapeutic options for KPLBS, we conducted an *in silico* molecular docking screen of FDA-approved drugs, followed by a functional APC screen to discover potential targeted therapeutics and to identify novel pharmacological tool molecules capable of probing wild-type and mutant channel function. Among the identified compounds, verapamil and lovastatin emerged as promising candidates. Verapamil was of particular interest due to its known inhibition of the *weaver* mutant (Lesage et al., 1995; Kofuji et al., 1996). While earlier reports suggested it was a poor wild-type channel blocker, our assay revealed robust block of hGIRK2 homomers at 30 µM (Lesage et al., 1995; Kofuji et al., 1996). However, its limited blood-brain barrier (BBB) penetration and status as a prominent P-glycoprotein substrate make it an unviable candidate for targeted *in vivo* inhibition of CNS hGIRK2 homomers. Furthermore, we found that lovastatin acutely blocked hGIRK2 currents. Statins are known to indirectly and chronically decrease mGIRK currents via cholesterol depletion (Deng et al., 2012; Bukiya et al., 2017, 2019). Our acute data, along with a recent computational study that found stable binding of rosuvastatin with GIRK2 at the PIP_2_ binding site, suggest that statins also possess the ability to directly inhibit the GIRK2 channel itself (Jeremic et al., 2025).

Pharmacological characterization of these two drugs revealed that the G154C mutation disrupts effective Gβγ coupling. In wild-type hGIRK2, eletriptan exhibited a Gβγ-dependent left-shift in potency, and nefazodone displayed highly cooperative, Gβγ-dependent block. Both dependencies were lost in the hGIRK2_G154C_ mutant. The loss of basal Gβγ activation, combined with the altered pharmacological profile, provides functional evidence that the SF mutation functionally uncouples the channel from its G-protein modulation.

Beyond demonstrating the loss of allosteric coupling, eletriptan’s biophysical profile allowed us to localize its site of action within the cytoplasmic pore. Notably, the predicted docking pose of eletriptan in the cytoplasmic vestibule (Figure 8) is consistent with previous studies showing that cationic blockers of inwardly rectifying potassium channels can occupy intracellularly accessible pore regions (De Boer et al., 2010; Takanari et al., 2013; Marmolejo-Murillo et al., 2017). Mechanistically, we found that eletriptan behaves as a classic intracellular pore blocker. At physiological pH, eletriptan is predominantly protonated, facilitating its access to the aqueous cytosol after crossing the plasma membrane (Tfelt-Hansen et al., 2000; Milton et al., 2002). However, eletriptan might be physically too bulky to pass through the narrow G-loop gate to reach the transmembrane (TM) inner vestibule (Hibino et al., 2010; Glaaser and Slesinger, 2015; Hines et al., 2017). Instead, the positively charged drug is drawn into the highly electronegative cytoplasmic pore, occluding the ion pathway just outside the Helix Bundle Crossing (HBC) gate (Nishida and MacKinnon, 2002; Hibino et al., 2010). This interpretation is further supported by the calculated fractional electrical distance (δ = 0.21 – 0.29), indicating that the drug sits shallowly in the pore, displacing local K⁺ ions outwards through the SF, which is thought to underlie the block’s apparent voltage sensitivity (Xu et al., 2009). The increased potency of eletriptan in the activated state suggests state-dependent inhibition, wherein Gβγ-induced conformational changes in the cytoplasmic domain likely enhance the drug’s binding affinity.

Eletriptan is an approved triptan used for the treatment of acute migraine (McCormack and Keating, 2006; Cameron et al., 2015). Despite potent channel block *in vitro*, an effect on CNS hGIRK2 in patients receiving eletriptan is unlikely, due to its limited brain penetrance (Svane et al., 2024). While eletriptan proved useful for confirming the mutant channel’s uncoupled state *in vitro*, it is an unviable therapeutic candidate for the treatment of KPLBS.

In addition, our APC screen also identified the atypical antidepressants vilazodone and nefazodone as potent hGIRK2 blockers (Plenge et al., 2021). Nefazodone is an atypical antidepressant of the serotonin antagonist and reuptake inhibitor (SARI) class (El-Kasaby et al., 2024). Given its higher potency and greater brain exposure at clinical doses, nefazodone was selected for detailed MPC analysis. Nefazodone is highly lipophilic and exists predominantly in its neutral, unionized state at physiological pH (Joshi et al., 1998).

In the wild-type + Gβγ condition, nefazodone block was virtually absent at low concentrations before showing strong inhibition at 3 µM, yielding a steep Hill coefficient of 3.13. Because the Hill coefficient represents a lower estimate for interacting sites, this suggests that multiple nefazodone molecules may contribute cooperatively to channel inhibition. Together with its high lipophilicity, these observations are consistent with a mechanism distinct from classical voltage-dependent pore block, although the precise binding site remains unresolved. Importantly, voltage dependence alone cannot unambiguously define blocker location or mechanism. Previous studies on Kir channels have demonstrated voltage-independent inhibition despite evidence for direct interactions within the cytoplasmic pore, suggesting that apparent voltage dependence may reflect not only blocker position but also channel gating and permeant-ion occupancy (Heginbotham and Kutluay, 2004; Koepple et al., 2017). Thus, while our data argue against a simple classical pore-blocking mechanism, they do not exclude pore-associated, allosteric, or other gating-coupled modes of inhibition (Shalomov et al., 2025). The strong cooperativity and marked dependence on Gβγ-mediated channel activation further suggest that channel conformation plays an important role in nefazodone action (Shalomov et al., 2025).

When nefazodone was assayed previously to explore its activity on GIRK1/2 and GIRK1/4 heteromers, even in the absence of Gβγ stimulation, it showed only partial inhibition of 35.9% and 30.3% at 100 µM (Kobayashi et al., 2011). Under our own experimental conditions without Gβγ overexpression, nefazodone achieved near-complete block of hGIRK2 homotetramers at 30 µM, a concentration well below the 100 µM dose required to elicit even weak, partial inhibition of heteromers (Kobayashi et al., 2011). If confirmed, such subtype selectivity could provide a useful pharmacological approach for probing GIRK2 homomer function in native systems.

Clinically achievable nefazodone concentrations may be sufficient to inhibit hGIRK2 homomers, particularly given its accumulation in the brain (Greene and Barbhaiya, 1997; Nacca et al., 1998). There is historical precedent targeting mGIRK2_G154S_ channels in the *weaver* mouse using the SSRI fluoxetine, successfully rescuing motor deficits and cerebellar cell death (Takahashi et al., 2006). However, classical antidepressants promiscuously block wild-type GIRK1/2, GIRK2, and GIRK1/4 channels, likely contributing to side effects such as seizure risk associated with fluoxetine (Borys et al., 1992; Kobayashi et al., 2004; Kolbeck et al., 2024). Nefazodone does not share the well-documented convulsant liability associated with fluoxetine (Davis et al., 1997; Benson et al., 2000). Whether this difference relates to distinct GIRK subtype selectivity profiles remains unknown.

It is therefore tempting to speculate that nefazodone could attenuate the pathological ion flux through the dysfunctional hGIRK2 homomer channels. Nefazodone’s established pharmacological profile provides a foundation for preclinical evaluation. However, any future therapeutic development must account for its rare but serious risk of hepatotoxicity (Stewart, 2002), which led to its withdrawal from several markets.

In summary, we demonstrate that the newly identified hGIRK2_G154C_ variant underlying a mild case of KPLBS causes severe biophysical defects *in vitro*, including a widened SF, loss of potassium selectivity, and uncoupling of the channel from the Gβγ complex. Through a combination of *in silico* and *in vitro* screens, we identified eletriptan and nefazodone as potent hGIRK2 homomer inhibitors. While eletriptan acts as a classic intracellular pore blocker, nefazodone functions as a highly cooperative, possibly subtype-selective inhibitor of hGIRK2 homomers under clinically relevant concentrations. Nefazodone thus represents both a promising pharmacological tool for probing native GIRK2 homomers and a potential candidate for further investigation in the context of KPLBS.

## Data Availability Statement

The raw electrophysiological data supporting the conclusions of this article will be made available by the authors, without undue reservation. The processed molecular dynamics datasets, trajectories, and rendering scripts presented in this study can be found in online repositories at: https://gitlab.com/vihuhol188/girk2_g156c/. The custom Python scripts used for the rundown correction of both the manual and automated patch-clamp data can be found at: https://github.com/MANetzer/patch-clamp-rundown-correction.

## Conflict of Interest

The authors declare that the research was conducted in the absence of any commercial or financial relationships that could be construed as a potential conflict of interest.

## Author Contributions

*Conceptualization*: MN, ND, ASW. *Methodology*: MN, IS, TF, ND, ASW. *Investigation*: MN, IS, ASW. *Formal analysis*: MN, IS, ND, ASW. *Supervision*: ND, ASW. *Writing – original draft*: MN, IS, ND, ASW. *Writing – review and editing*: all. *Funding acquisition*: ASW, ND.

## Funding

This work was supported by the doctoral program “Molecular drug targets” W1232 (M.N, T.F. and A.S.-W.), the DOC fellowship 26156 of the Austrian Academy of Sciences (ÖAW) (T.F.), the Israel Science Foundation (ISF) grant #581_2022 (N.D.).

## Acknowledgements

The computational results have been achieved using the Austrian Scientific Computing (ASC) infrastructure. The authors gratefully acknowledge Univ. Prof. Dr. Steffen Hering for his support and provision of laboratory facilities, and thank the entire team at ChanPharm GmbH for their assistance.

## Supplementary material

**Supplementary Figure 1.**
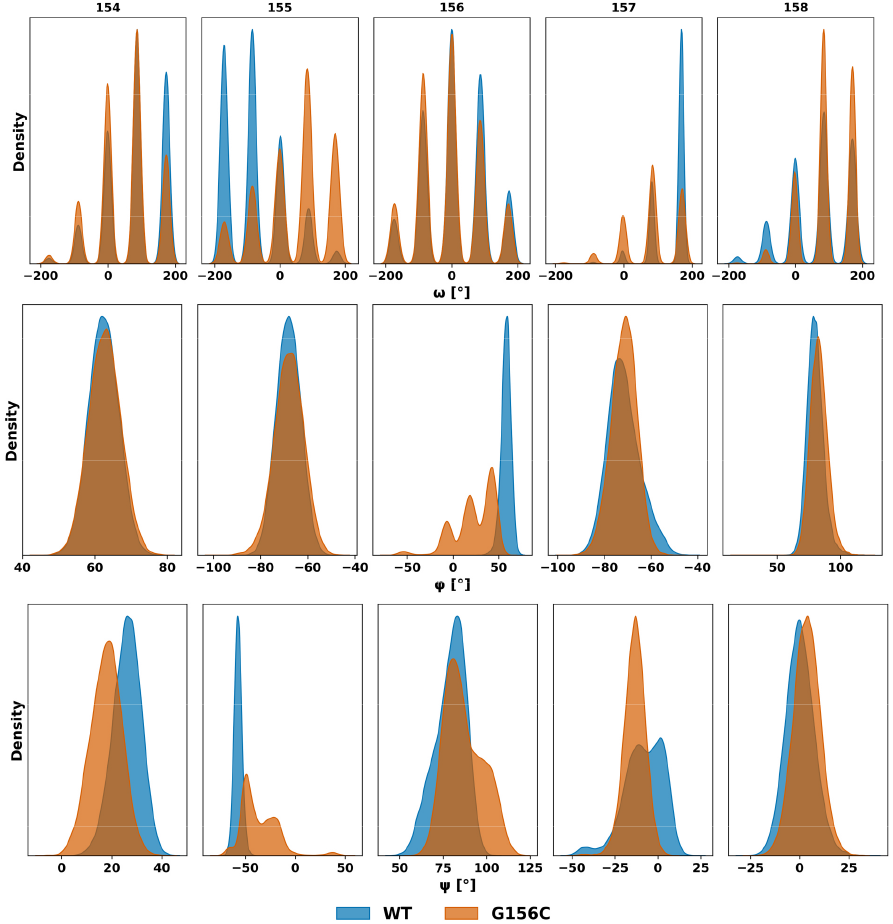
Backbone dihedral angle distributions of selectivity filter residues (154–158) in mGIRK2_WT_ and mGIRK2_G156C_ during MD simulations. Distributions of ω, φ, and ψ angles are shown for each residue, highlighting substantial conformational changes induced by the G156C mutation. Data represents aggregate structural dynamics sampled over 1 µs simulations.

**Supplementary Figure 2.**
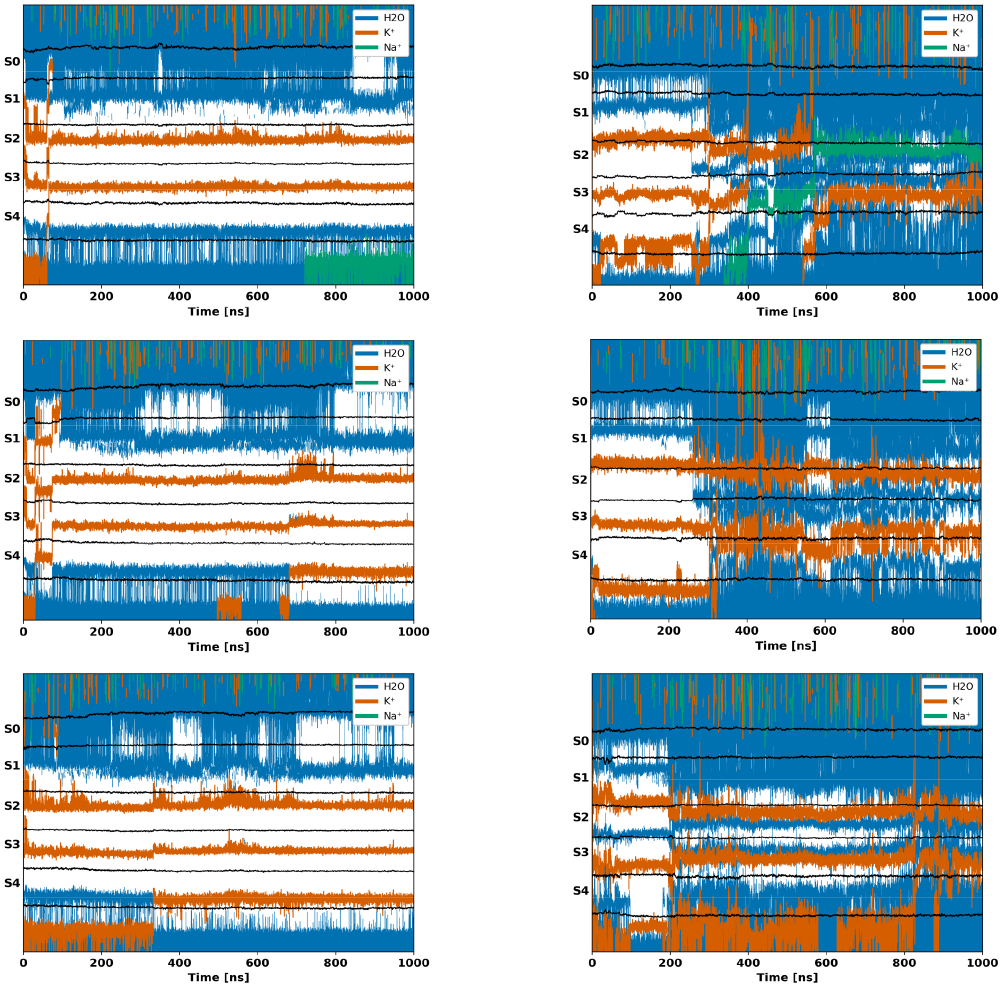
Ion and water occupancy in the selectivity filter of mGIRK2_WT_ and mGIRK2_G156C_ across independent MD simulation replicas. Time-resolved occupancy of K^+^, Na^+^, and H_2_0 at sites S0–S4 over 1 µs simulations is shown for three independent replicas, confirming the reproducibility of aberrant ion permeation and structural deformation in the mutant channel.

**Supplementary Figure 3.**
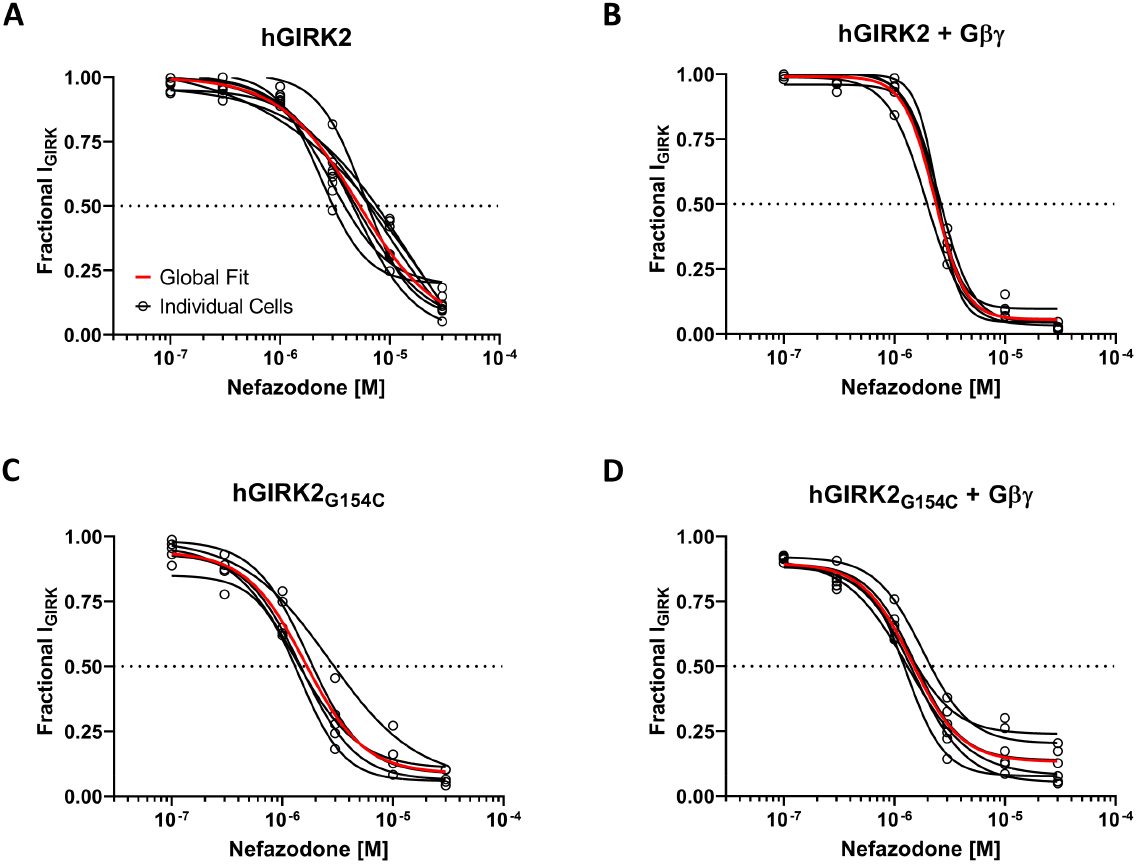
Individual cell dose-response curve fits for nefazodone using MPC. **(A) – (D)** Dose-response curves of individual cells of wild-type hGIRK2 and mutant G154C channels in the presence and absence of co-overexpressed Gβγ. Currents were measured at –120 mV in 48 mM external K+ solutions and normalized to 3 mM Ba2+ block for wild-type conditions or maximum negative amplitude for mutant conditions.

**Supplementary Figure 4.**
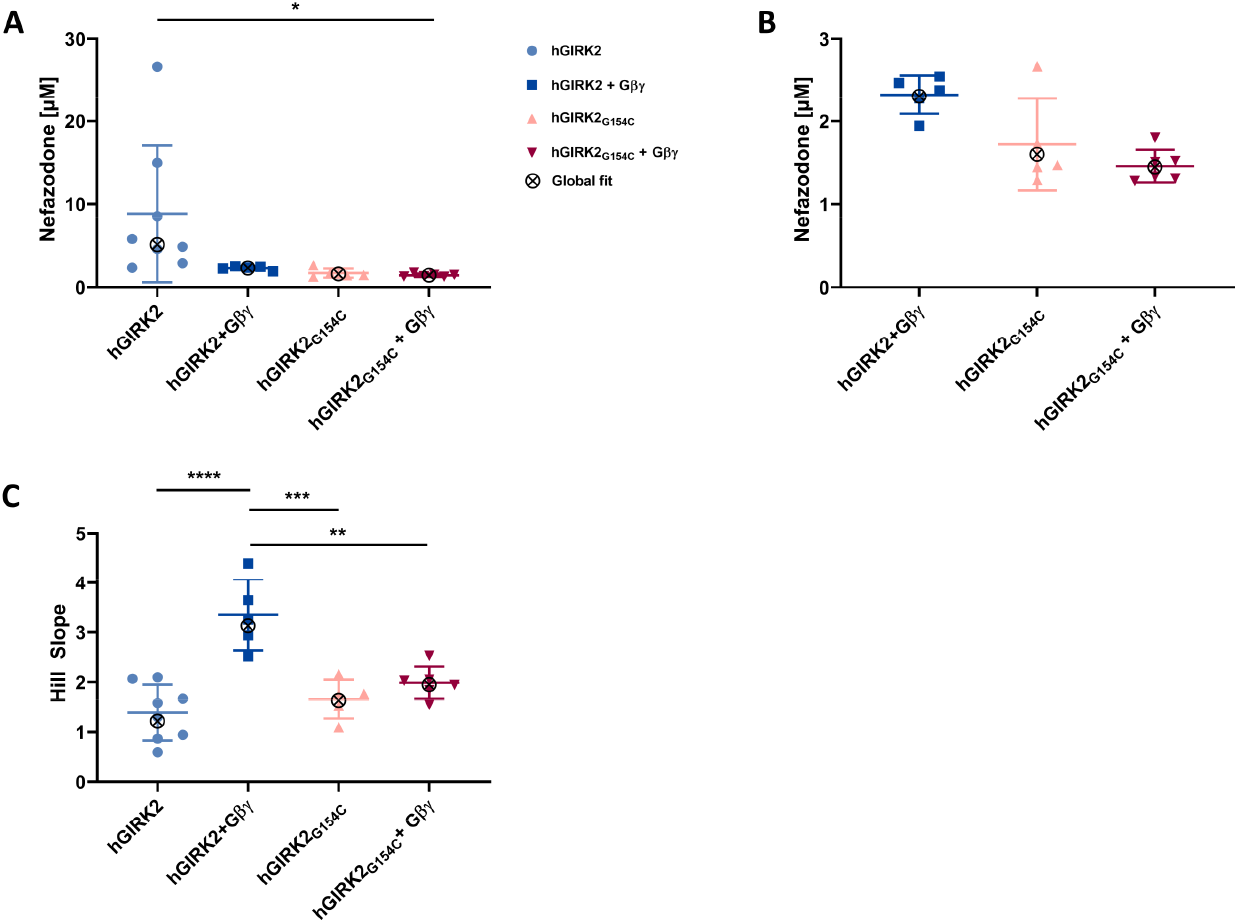
Individual cell Hill slopes and IC_50_ values of DR curve fits for nefazodone using MPC. Values were extracted from curve fits seen in Supplementary Figure 3. **(A)** depicts IC_50_ values of different test conditions, with global fit values overlaid. **(B)** shows the same data as (A) but without hGIRK2. **(C)** similarly shows Hill slope values extracted from curve fits seen in Supplementary Figure 3, global fit values are overlaid as well. Statistical significance was determined using ordinary one-way ANOVA followed by Tukey’s post hoc test. The different levels of significance are indicated as follows: ** *p* < .01, *** *p* < .001, **** *p* < .0001.

**Supplementary Figure 5.**
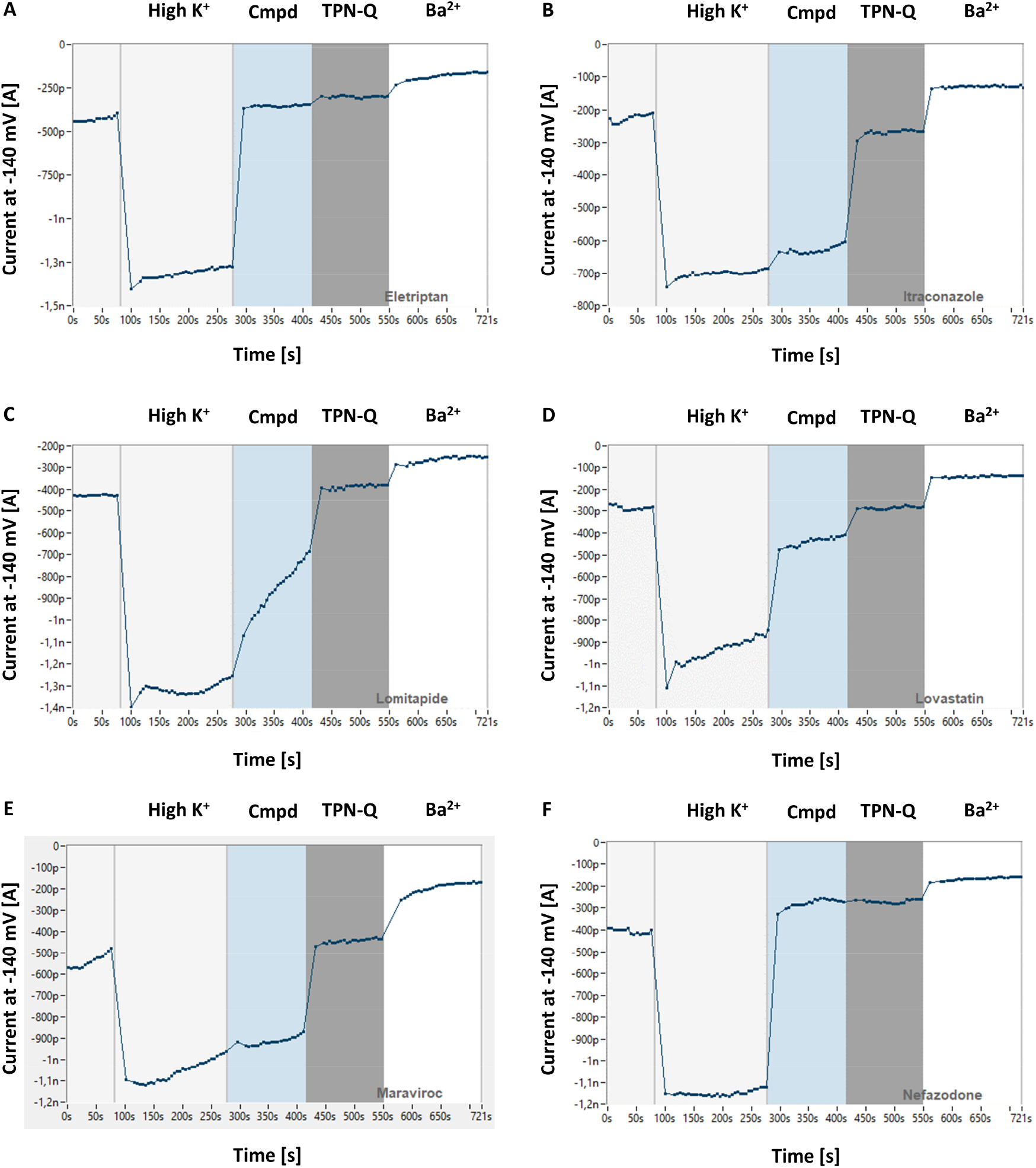
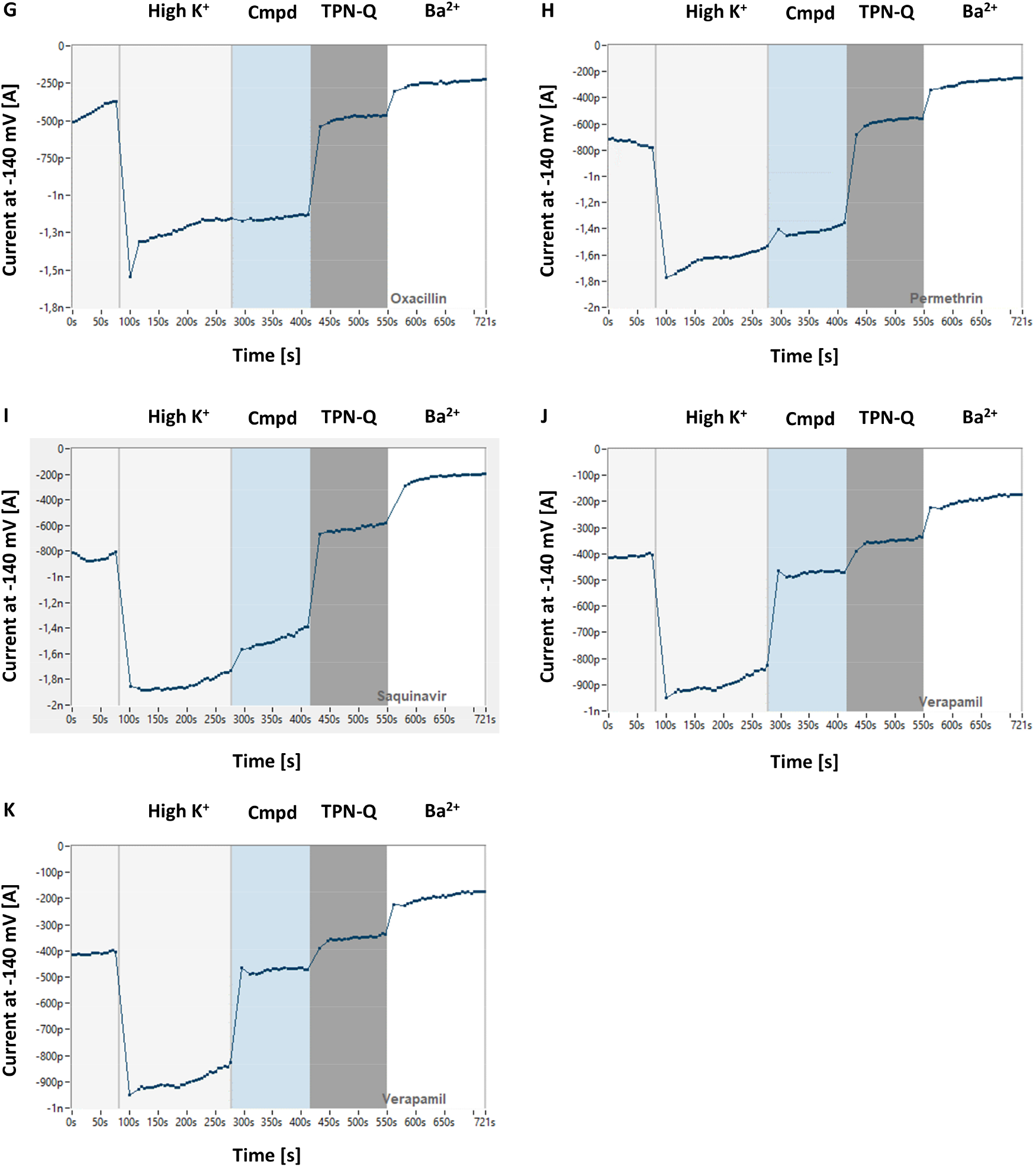
Representative automated patch clamp recordings of all tested compounds. **(A) – (K)** Average currents over time of CHO cells expressing hGIRK2 + Gβγ treated sequentially with 30 µM of test compound, 400 nM TPN-Q, and 3 mM Ba^2+^. Only cells passing rigorous quality control criteria (series resistance <20 MΩ, capacitance <40 pF, membrane resistance >100 MΩ) were included. Panels depict results from one representative experimental run of eletriptan, itraconazole, lomitapide, lovastatin, maraviroc, nefazodone, oxacillin, permethrin, saquinavir, verapamil, and vilazodone, respectively, corresponding to (A) – (K).

**Supplementary Figure 6.**
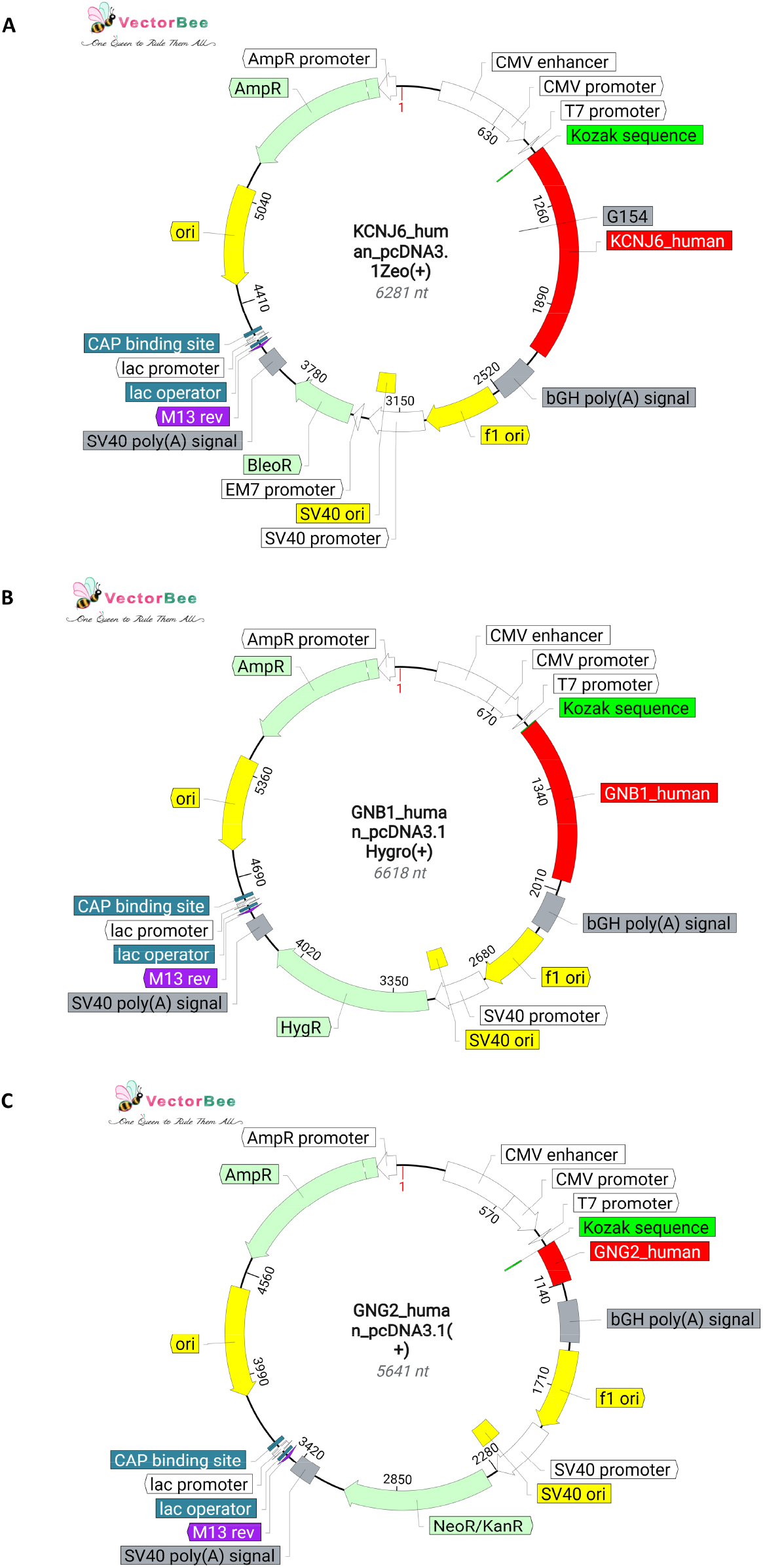
DNA plasmid maps of constructs used in this study. **(A) – (C)** Detailed vector maps illustrating the pcDNA3.1(+) mammalian expression constructs containing the human *KCNJ6* (with the G154C mutation site highlighted), *GNB1*, and *GNG2* genes. Sequences were verified via standard Sanger sequencing, and plasmid maps were created with VectorBee (VectorBuilder).

---

**Supplementary Table 1:** Changes in the force constant of a protein during equilibration of the system.

| Stage | Force constant |  |
| --- | --- | --- |
|  | BB, kJ·mol<sup>-1</sup> nm<sup>-2</sup> | SC, kJ·mol<sup>-1</sup> nm<sup>-2</sup> |
| NVT | 4000 | 2000 |
| NVT | 2000 | 1000 |
| NPT | 1000 | 500 |
| NPT | 500 | 200 |
| NPT | 200 | 50 |
| NPT | 50 | 0 |

*Note.* Position restraint force constants applied to the protein backbone (BB) and side chains (SC) across the six stages of system equilibration. NVT denotes the canonical ensemble (constant number, volume, and temperature) and NPT denotes the isothermal-isobaric ensemble (constant number, pressure, and temperature).

**Supplementary Table 2:**
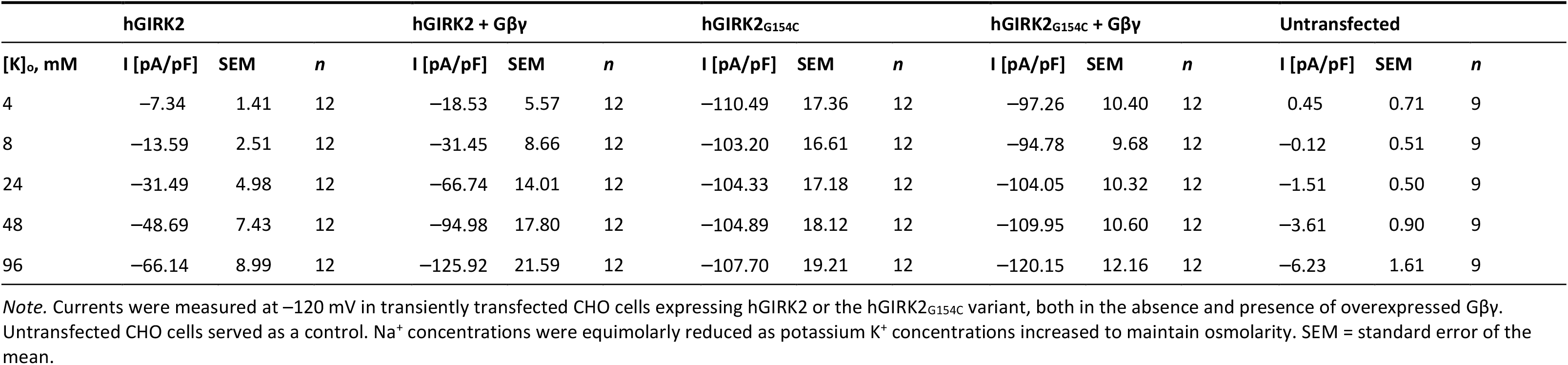
Maximal Inward Current Densities Across Varying Extracellular Potassium Concentrations.

**Supplementary Table 3:** Reversal Potentials Across Varying Extracellular Potassium Concentrations.

| [K]<sub>o</sub>, mM | hGIRK2 |  |  | hGIRK2 + Gβγ |  |  | hGIRK2<sub>G154C</sub> |  |  | hGIRK2<sub>G154C</sub> + Gβγ |  |  |
| --- | --- | --- | --- | --- | --- | --- | --- | --- | --- | --- | --- | --- |
|  | V<sub>rev</sub> [mV] | SEM | n | V<sub>rev</sub> [mV] | SEM | n | V<sub>rev</sub> [mV] | SEM | n | V<sub>rev</sub> [mV] | SEM | n |
| 4 | -89.04 | 2.60 | 9 | -89.36 | 1.56 | 8 | -6.34 | 1.91 | 8 | -14.64 | 2.91 | 9 |
| 8 | -71.90 | 1.29 | 11 | -73.74 | 2.19 | 9 | -5.73 | 1.33 | 9 | -16.45 | 2.68 | 11 |
| 24 | -44.07 | 0.93 | 11 | -46.75 | 1.56 | 9 | -5.51 | 1.41 | 9 | -12.34 | 2.37 | 11 |
| 48 | -27.24 | 1.02 | 11 | -27.77 | 2.42 | 10 | -4.17 | 1.46 | 11 | -9.57 | 1.32 | 12 |
| 96 | -9.38 | 0.74 | 11 | -10.60 | 2.21 | 9 | -1.97 | 1.86 | 12 | -4.83 | 1.43 | 12 |

*Note.* Reversal potentials (V<sub>rev</sub>) were measured in transiently transfected CHO cells after the subtraction of Ba<sup>2+</sup>-sensitive currents. Na<sup>+</sup> concentrations were equimolarly reduced as potassium K<sup>+</sup> concentrations increased. SEM = standard error of the mean.

**Supplementary Table 4:** Relative Permeability Ratios of Na^+^ and K^+^.

|  | **p<sub>Na</sub>/p<sub>K</sub>** |  |  |
| --- | --- | --- | --- |
|  | **Ratio** | **SEM** | **n** |
| hGIRK2 | 0.0027 | 0.0050 | 11 |
| hGIRK2 + Gβγ | 0.0120 | 0.0162 | 8 |
| hGIRK2<sub>G154C</sub> | 0.7464 | 0.0405 | 9 |
| hGIRK2<sub>G154C</sub> + Gβγ | 0.6026 | 0.0467 | 11 |

*Note.* The relative permeability ratio (p<sub>Na</sub>/p<sub>K</sub>) was determined according to the Goldman-Hodgkin-Katz voltage equation. Extracellular solutions of 8 mM K<sup>+</sup> (with 136 mM Na<sup>+</sup>) and 96 mM K<sup>+</sup> (with 48 mM Na<sup>+</sup>) were used to calculate the shift in V<sub>rev</sub>. SEM = standard error of the mean.

**Supplementary Table 5:** Statistical Analysis of Permeability Ratios.

|  | **Summary** | **p value** |
| --- | --- | --- |
| hGIRK2 vs. hGIRK2 + Gβγ | ns | 0.9449 |
| hGIRK2 vs. hGIRK2<sub>G154C</sub> | **** | <0.0001 |
| hGIRK2 vs. hGIRK2<sub>G154C</sub> + Gβγ | **** | <0.0001 |
| hGIRK2 + Gβγ vs. hGIRK2<sub>G154C</sub> | **** | <0.0001 |
| hGIRK2 + Gβγ vs. hGIRK2<sub>G154C</sub> + Gβγ | **** | <0.0001 |
| hGIRK2<sub>G154C</sub> vs. hGIRK2<sub>G154C</sub> + Gβγ | ns | 0.1283 |

*Note.* Statistical significance between conditions was determined using the Brown-Forsythe and Welch ANOVA, followed by the Games-Howell post hoc multiple comparisons test. ns = not significant. \*\*\*\* *p* < .0001

**Supplementary Table 6:** Inward Rectification Indices for Wild-Type and Mutant hGIRK2 Channels.

| **F<sub>ir</sub>** |  |  |  |
| --- | --- | --- | --- |
|  | **Ratio** | **SEM** | ***n*** |
| hGIRK2 | 0.1855 | 0.0135 | 11 |
| hGIRK2 + Gβγ | 0.2521 | 0.0350 | 9 |
| hGIRK2<sub>G154C</sub> | 0.5799 | 0.0403 | 12 |
| hGIRK2<sub>G154C</sub> + Gβγ | 0.6524 | 0.0336 | 12 |

*Note.* The inward rectification index (F<sub>ir</sub>) was calculated by dividing the current at 40 mV positive of V<sub>rev</sub> by the current at 40 mV negative V<sub>rev</sub>. SEM = standard error of the mean.

**Supplementary Table 7:**
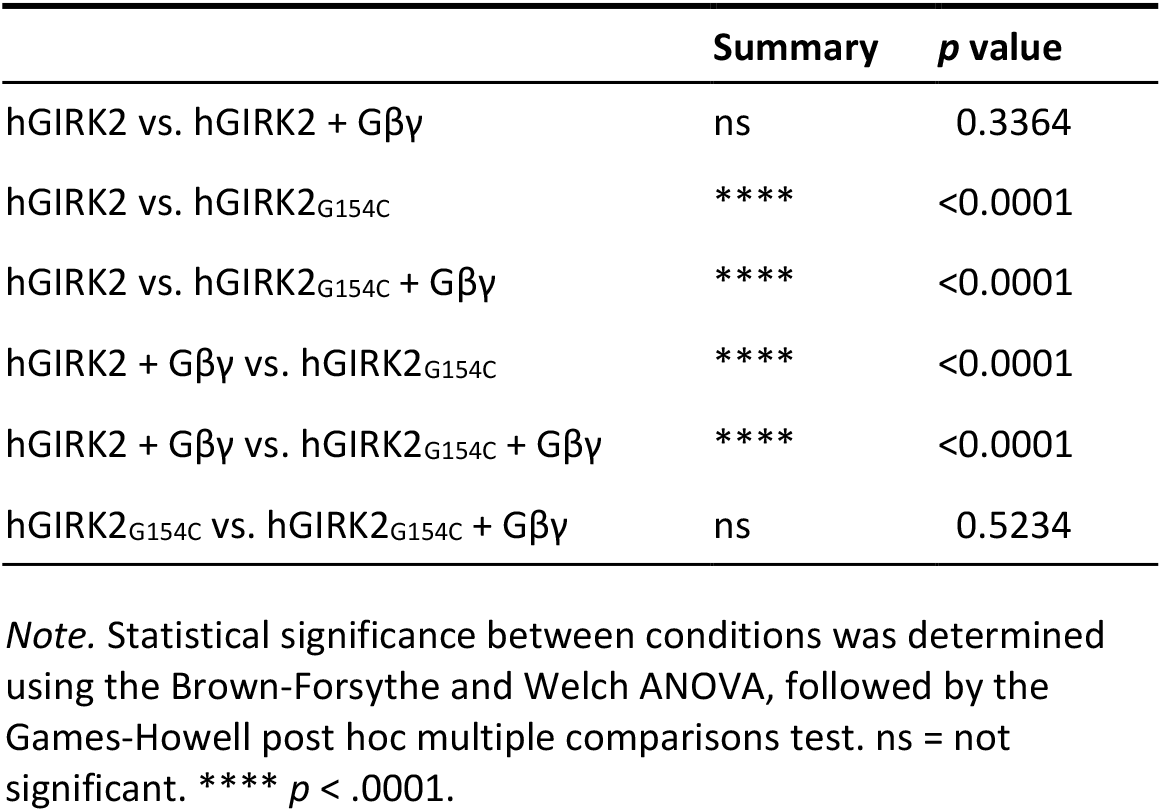
Statistical Analysis of Inward Rectification Indices.

|  | **Summary** | ***p* value** |
| --- | --- | --- |
| hGIRK2 vs. hGIRK2 + Gβγ | ns | 0.3364 |
| hGIRK2 vs. hGIRK2<sub>G154C</sub> | **** | <0.0001 |
| hGIRK2 vs. hGIRK2<sub>G154C</sub> + Gβγ | **** | <0.0001 |
| hGIRK2 + Gβγ vs. hGIRK2<sub>G154C</sub> | **** | <0.0001 |
| hGIRK2 + Gβγ vs. hGIRK2<sub>G154C</sub> + Gβγ | **** | <0.0001 |
| hGIRK2<sub>G154C</sub> vs. hGIRK2<sub>G154C</sub> + Gβγ | ns | 0.5234 |

*Note.* Statistical significance between conditions was determined using the Brown-Forsythe and Welch ANOVA, followed by the Games-Howell post hoc multiple comparisons test. ns = not significant. \*\*\*\* *p* < .0001.

**Supplementary Table 8:**
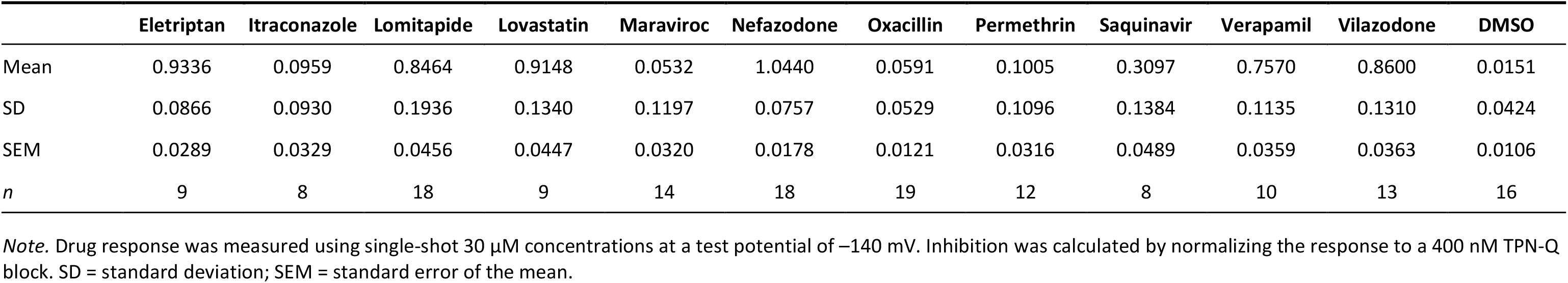
Automated Patch Clamp Assay Inhibition by Selected Compounds at 30 µM.

|  | Eletriptan | Itraconazole | Lomitapide | Lovastatin | Maraviroc | Nefazodone | Oxacillin | Permethrin | Saquinavir | Verapamil | Vilazodone | DMSO |
| --- | --- | --- | --- | --- | --- | --- | --- | --- | --- | --- | --- | --- |
| Mean | 0.9336 | 0.0959 | 0.8464 | 0.9148 | 0.0532 | 1.0440 | 0.0591 | 0.1005 | 0.3097 | 0.7570 | 0.8600 | 0.0151 |
| SD | 0.0866 | 0.0930 | 0.1936 | 0.1340 | 0.1197 | 0.0757 | 0.0529 | 0.1096 | 0.1384 | 0.1135 | 0.1310 | 0.0424 |
| SEM | 0.0289 | 0.0329 | 0.0456 | 0.0447 | 0.0320 | 0.0178 | 0.0121 | 0.0316 | 0.0489 | 0.0359 | 0.0363 | 0.0106 |
| *n* | 9 | 8 | 18 | 9 | 14 | 18 | 19 | 12 | 8 | 10 | 13 | 16 |

*Note.* Drug response was measured using single-shot 30 μM concentrations at a test potential of $-140$ mV. Inhibition was calculated by normalizing the response to a 400 nM TPN-Q block. SD = standard deviation; SEM = standard error of the mean.

**Supplementary Table 9:** Automated Patch Clamp Assay Inhibition for Eletriptan and Nefazodone Across Multiple Concentrations.

| hGIRK2 + Gβ $\gamma$ | | | | hGIRK2 + Gβ $\gamma$ | | | |
| --- | --- | --- | --- | --- | --- | --- | --- |
| Eletriptan [μM] | Mean | SD | *n* | Nefazodone [μM] | Mean | SD | *n* |
| 0.1 | 0.0402 | 0.0671 | 3 | 0.1 | 0.0040 | 0.0815 | 4 |
| 0.3 | 0.0991 | 0.0759 | 8 | 0.3 | 0.0290 | 0.0180 | 3 |
| 1 | 0.2881 | 0.0733 | 3 | 1 | 0.0185 | 0.0257 | 3 |
| 3 | 0.5672 | 0.0563 | 4 | 3 | 0.5137 | 0.1854 | 3 |
| 10 | 0.7878 | 0.0591 | 6 | 10 | 0.8477 | 0.0986 | 7 |
| 30 | 0.9804 | 0.1179 | 5 | 30 | 0.9993 | 0.0181 | 6 |

*Note.* Drug response was measured at a test potential of $-140$ mV and normalized to a 400 nM TPN-Q block using automated patch clamp. SD = standard deviation.

**Supplementary Table 10:**
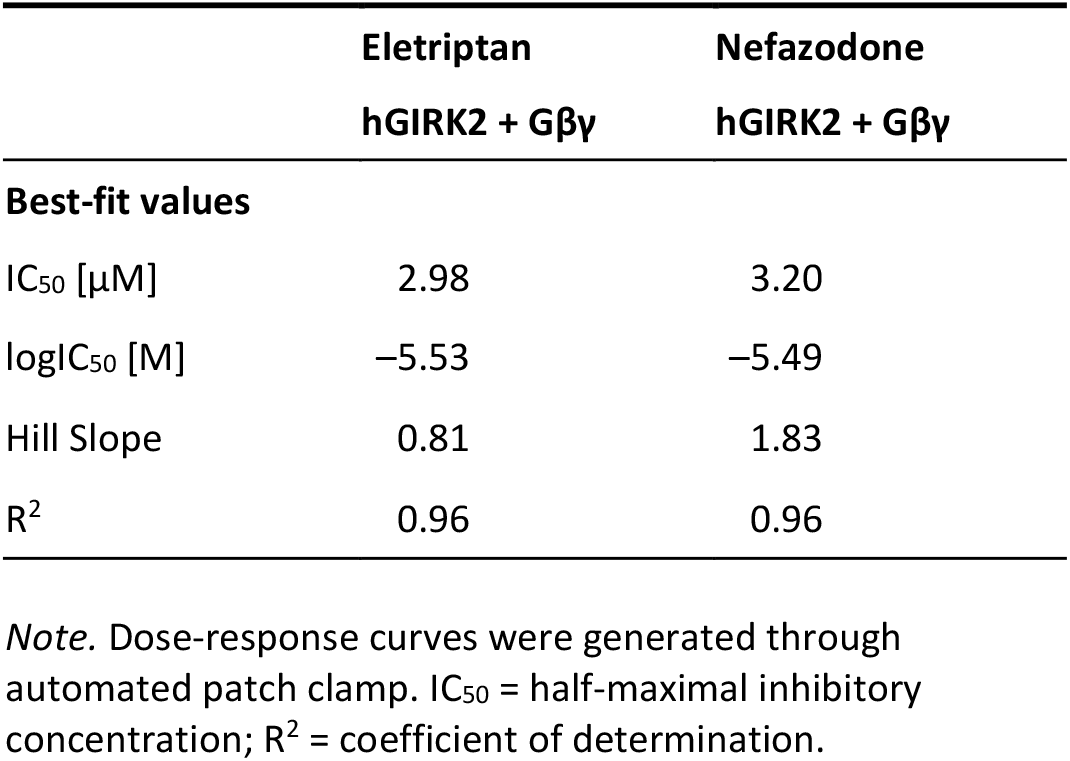
Best-Fit Values for Eletriptan and Nefazodone Dose-Response Curves Using Automated Patch Clamp.

|  | Eletriptan | Nefazodone |
| --- | --- | --- |
|  | hGIRK2 + Gβγ | hGIRK2 + Gβγ |
| **Best-fit values** |  |  |
| IC<sub>50</sub> [μM] | 2.98 | 3.20 |
| logIC<sub>50</sub> [M] | −5.53 | −5.49 |
| Hill Slope | 0.81 | 1.83 |
| R<sup>2</sup> | 0.96 | 0.96 |

*Note.* Dose-response curves were generated through automated patch clamp. IC<sub>50</sub> = half-maximal inhibitory concentration; R<sup>2</sup> = coefficient of determination.

**Supplementary Table 11:** Best-Fit Values for Nefazodone Dose-Response Curves Using Manual Patch Clamp.

|  | Nefazodone |  |  |  |
| --- | --- | --- | --- | --- |
|  | hGIRK2 | hGIRK2 + Gβγ | hGIRK2<sub>G154C</sub> | hGIRK2<sub>G154C</sub> + Gβγ |
| **Best-fit values** |  |  |  |  |
| IC<sub>50</sub> [μM] | 5.12 | 2.30 | 1.60 | 1.45 |
| logIC<sub>50</sub> [M] | −5.32 | −5.64 | −5.80 | −5.84 |
| Hill Slope | 1.22 | 3.13 | 1.63 | 1.95 |
| R<sup>2</sup> | 0.97 | 0.99 | 0.97 | 0.96 |

*Note.* Dose-response curves were generated through manual patch clamp. IC<sub>50</sub> = half-maximal inhibitory concentration; R<sup>2</sup> = coefficient of determination.

**Supplementary Table 12:** Manual Patch Clamp Assay Inhibition for Nefazodone Across Multiple Concentrations.

| Nefazodone [μM] | hGIRK2 | | | hGIRK2 + Gβy | | | hGIRK2<sub>G154C</sub> | | | hGIRK2<sub>G154C</sub> + Gβy | | |
| --- | --- | --- | --- | --- | --- | --- | --- | --- | --- | --- | --- | --- |
|  | Mean | SD | *n* | Mean | SD | *n* | Mean | SD | *n* | Mean | SD | *n* |
| 0.1 | 0.0211 | 0.0264 | 8 | 0.0000 | 0.0166 | 5 | 0.0535 | 0.0385 | 5 | 0.0847 | 0.0099 | 6 |
| 0.3 | 0.0355 | 0.0377 | 8 | 0.0237 | 0.0326 | 5 | 0.1321 | 0.0565 | 5 | 0.1679 | 0.0393 | 6 |
| 1 | 0.0721 | 0.0375 | 8 | 0.0670 | 0.0538 | 5 | 0.3128 | 0.0774 | 5 | 0.3403 | 0.0563 | 6 |
| 3 | 0.3749 | 0.0966 | 8 | 0.6624 | 0.0502 | 5 | 0.7055 | 0.1022 | 5 | 0.7342 | 0.0828 | 6 |
| 10 | 0.6515 | 0.0773 | 8 | 0.9045 | 0.0349 | 5 | 0.8388 | 0.0699 | 5 | 0.8205 | 0.0847 | 6 |
| 30 | 0.8868 | 0.0392 | 8 | 0.9731 | 0.0120 | 5 | 0.9263 | 0.0280 | 5 | 0.8854 | 0.0643 | 6 |

*Note.* Drug response was measured at a test potential of $-120$ mV using manual patch clamp. For hGIRK2 wild-type conditions, responses were normalized to a 3 mM Ba<sup>2+</sup> block. For hGIRK2<sub>G154C</sub> mutant conditions, responses were normalized to the maximal negative amplitude. SD = standard deviation.

**Supplementary Table 13:** Statistical Analysis of Manual Patch Clamp Nefazodone IC_50_ Values On Individual Fit Basis.

|  | Summary | *p* value |
| --- | --- | --- |
| hGIRK2 vs. hGIRK2 + Gβy | **** | <0.0001 |
| hGIRK2 vs. hGIRK2<sub>G154C</sub> | ns | 0.7975 |
| hGIRK2 vs. hGIRK2<sub>G154C</sub> + Gβy | ns | 0.1743 |
| hGIRK2 + Gβy vs. hGIRK2<sub>G154C</sub> | *** | 0.0003 |
| hGIRK2 + Gβy vs. hGIRK2<sub>G154C</sub> + Gβy | ** | 0.0017 |
| hGIRK2<sub>G154C</sub> vs. hGIRK2<sub>G154C</sub> + Gβy | ns | 0.7232 |

*Note.* Statistical significance between conditions was determined using Ordinary One-Way ANOVA, followed by the Tukey post hoc multiple comparisons test. ns = not significant. \*\* $p < .01$ , \*\*\* $p < .001$ , \*\*\*\* $p < .0001$ .

**Supplementary Table 14:** Statistical Analysis of Manual Patch Clamp Nefazodone Hill Slope Values On Individual Fit Basis.

|  | Summary | *p* value |
| --- | --- | --- |
| hGIRK2 vs. hGIRK2 + Gβy | ns | 0.1218 |
| hGIRK2 vs. hGIRK2<sub>G154C</sub> | ns | 0.0808 |
| hGIRK2 vs. hGIRK2<sub>G154C</sub> + Gβy | * | 0.0498 |
| hGIRK2 + Gβy vs. hGIRK2<sub>G154C</sub> | ns | 0.9973 |
| hGIRK2 + Gβy vs. hGIRK2<sub>G154C</sub> + Gβy | ns | 0.9910 |
| hGIRK2<sub>G154C</sub> vs. hGIRK2<sub>G154C</sub> + Gβy | ns | 0.9997 |

*Note.* Statistical significance between conditions was determined using Ordinary One-Way ANOVA, followed by the Tukey post hoc multiple comparisons test. ns = not significant. \* $p < .05$ .

**Supplementary Table 15:** Voltage-Dependence of Nefazodone Block Measured by Manual Patch Clamp.

| hGIRK2 |  |  |  | hGIRK2 + Gβγ |  |  | hGIRK2<sub>G154C</sub> |  |  | hGIRK2<sub>G154C</sub> + Gβγ |  |  |
| --- | --- | --- | --- | --- | --- | --- | --- | --- | --- | --- | --- | --- |
| Nefazodone [μM] | Slope | Summary | p-value | Slope | Summary | p-value | Slope | Summary | p-value | Slope | Summary | p-value |
| 0.1 | 0.0003 | ns | 0.3786 | 0.0001 | ns | 0.5643 | −0.0001 | ns | 0.6563 | −0.0001 | ns | 0.4372 |
| 0.3 | 0.0007 | ns | 0.1892 | −0.0006 | ns | 0.3080 | −0.0002 | ns | 0.5254 | −0.0002 | ns | 0.2445 |
| 1 | 0.0013 | * | 0.0231 | 0.0008 | ns | 0.3622 | −0.0005 | ns | 0.2142 | −0.0001 | ns | 0.7264 |
| 3 | 0.0006 | ns | 0.0652 | 0.0002 | ns | 0.6253 | −0.0003 | ns | 0.5377 | 0.0000 | ns | 0.9649 |
| 10 | 0.0007 | ns | 0.0710 | 0.0002 | ns | 0.6287 | −0.0001 | ns | 0.8710 | 0.0001 | ns | 0.8373 |
| 30 | 0.0004 | ns | 0.2554 | 0.0003 | ns | 0.1034 | −0.0001 | ns | 0.7068 | 0.0000 | ns | 0.9162 |

Note. Data were transformed and fitted using simple linear regression at test voltages ranging from −115 mV to −35 mV. Slopes significantly different from zero indicate voltage-dependent block. ns = not significant. \* *p* < .05.

**Supplementary Table 16:** Best-Fit Values for Eletriptan Dose-Response Curves Using Manual Patch Clamp.

| Eletriptan |  |  |  |  |
| --- | --- | --- | --- | --- |
|  | hGIRK2 | hGIRK2 + Gβγ | hGIRK2<sub>G154C</sub> | hGIRK2<sub>G154C</sub> + Gβγ |
| Best-fit values |  |  |  |  |
| IC<sub>50</sub> [μM] | 2.88 | 1.49 | 3.09 | 3.68 |
| logIC<sub>50</sub> [M] | −5.54 | −5.83 | −5.51 | −5.43 |
| Hill Slope | 0.77 | 0.78 | 0.84 | 1.07 |
| R<sup>2</sup> | 0.96 | 0.96 | 0.96 | 0.98 |

Note. Dose-response curves were generated through manual patch clamp. IC<sub>50</sub> = half-maximal inhibitory concentration; R<sup>2</sup> = coefficient of determination.

**Supplementary Table 17:** Manual Patch Clamp Assay Inhibition for Eletriptan Across Multiple Concentrations.

| Eletriptan [μM] | hGIRK2 | | | hGIRK2 + Gβy | | | hGIRK2<sub>G154C</sub> | | | hGIRK2<sub>G154C</sub> + Gβy | | |
| --- | --- | --- | --- | --- | --- | --- | --- | --- | --- | --- | --- | --- |
|  | Mean | SD | *n* | Mean | SD | *n* | Mean | SD | *n* | Mean | SD | *n* |
| 0.3 | 0.1383 | 0.0642 | 5 | 0.1701 | 0.0477 | 6 | 0.0884 | 0.0525 | 5 | 0.0321 | 0.0400 | 5 |
| 1 | 0.2776 | 0.0590 | 6 | 0.3863 | 0.0697 | 6 | 0.2350 | 0.0654 | 5 | 0.1472 | 0.0489 | 5 |
| 3 | 0.4878 | 0.0625 | 6 | 0.5945 | 0.0868 | 6 | 0.4391 | 0.0345 | 6 | 0.3978 | 0.0672 | 5 |
| 10 | 0.6830 | 0.0532 | 6 | 0.7910 | 0.0866 | 6 | 0.6605 | 0.0589 | 5 | 0.6635 | 0.0551 | 5 |
| 30 | 0.8105 | 0.0638 | 6 | 0.9015 | 0.0381 | 5 | 0.7957 | 0.0635 | 6 | 0.8245 | 0.0283 | 5 |
| 100 | 0.8962 | 0.0480 | 6 | 0.9405 | 0.0350 | 6 |  |  |  |  |  |  |

*Note.* Drug response was measured at a test potential of $-120$ mV using manual patch clamp. For hGIRK2 wild-type conditions, responses were normalized to a 3 mM Ba<sup>2+</sup> block. For hGIRK2<sub>G154C</sub> mutant conditions, responses were normalized to a 1 mM QX-314 block. SD = standard deviation.

**Supplementary Table 18:** Voltage-Dependence of Eletriptan Block Measured by Manual Patch Clamp.

| Eletriptan [μM] | hGIRK2 | | | hGIRK2 + Gβy | | | hGIRK2<sub>G154C</sub> | | | hGIRK2<sub>G154C</sub> + Gβy | | |
| --- | --- | --- | --- | --- | --- | --- | --- | --- | --- | --- | --- | --- |
|  | Slope | Summary | *p*-value | Slope | Summary | *p*-value | Slope | Summary | *p*-value | Slope | Summary | *p*-value |
| 0.3 | -0.0012 | ns | 0.0792 | -0.0013 | * | 0.0050 | -0.0012 | * | 0.0012 | -0.0005 | ns | 0.1812 |
| 1 | -0.0013 | * | 0.0036 | -0.0023 | ** | 0.0009 | -0.0019 | **** | <0.0001 | -0.0015 | * | 0.0015 |
| 3 | -0.0025 | **** | <0.0001 | -0.0020 | ns | 0.0582 | -0.0023 | **** | <0.0001 | -0.0020 | ** | 0.0009 |
| 10 | -0.0018 | ** | 0.0005 | -0.0014 | * | 0.0449 | -0.0018 | *** | 0.0003 | -0.0015 | * | 0.0018 |
| 30 | -0.0007 | ns | 0.1004 | -0.0004 | ns | 0.2068 | -0.0010 | * | 0.0244 | -0.0007 | * | 0.0052 |

*Note.* Data were transformed and fitted using simple linear regression at test voltages ranging from $-115$ mV to $-35$ mV. Slopes significantly different from zero indicate voltage-dependent block. ns = not significant. \* $p < .05$ , $p < .01$ , \*\*\* $p < .001$ , \*\*\*\* $p < .0001$ .

## Notes

https://gitlab.com/vihuhol188/girk2_g156c/

https://github.com/MANetzer/patch-clamp-rundown-correction

### Competing Interest Statement

The authors have declared no competing interest.

